# In-House Manufacturing of 3D Culture Chips via Vacuum Thermoforming for Enhanced Imaging Applications

**DOI:** 10.1101/2025.02.26.640460

**Authors:** Aleksandra Fomina, Adam Tam, Jennifer Lam, Nila C. Wu, Nancy T. Li, Alison P. McGuigan

## Abstract

Engineered 3D in vitro cancer models, particularly those that facilitate image-based readouts capable of distinguishing the behavior of different cell populations, have become crucial tools in the discovery process. One such model, 96-GLAnCE (Gels for Live Analysis of Compartmentalized Environments), allows for longitudinal imaging of tumor cell dynamics and therapy response. However, the widespread adoption of 96-GLAnCE has been limited by the need for expensive, specialized fabrication equipment. To overcome this challenge, we have optimized a desktop vacuum thermoforming technique for the in-house production of 96- GLAnCE bottom chips using thin polystyrene films. This optimization has led to the reliable and consistent fabrication of devices. Notably, using thin polystyrene films reduces the overall thickness of the chips, enabling high-magnification imaging for studying primary tumor cell phenotypes in 3D with single-cell and subcellular resolution. Our thermoformed devices offer a flexible, cost-effective solution for addressing a wide range of biological questions across various time scales.

## 1. Introduction

Cancer is a complex, multifaceted disease that lacks effective therapies and has a poorly understood pathobiology(1,2). Ongoing research focuses on identifying disease progression mechanisms in gold-standard preclinical models, such as cancer cell lines cultured on 2D plastic and animal models. These preclinical models provide essential tools to study cancer cell and tumor biology; however, they have several disadvantages for the discovery process. 2D cancer cell line cultures lack tumor dimensionality, protein matrix, and other cell components. Animal models are considered more inefficient due to their high maintenance cost, time resources, and interspecies differences. Additionally, researchers have less control over the composition of the tumor microenvironment in animal models, which renders specific biological questions more challenging to pursue(3,4).

With the rise of the bioengineering field, physiologically representative 3D *in vitro* cancer models have emerged to complement findings in 2D systems and animal models and facilitate the discovery process. Several 3D *in vitro* cancer models exist, including tumor spheroids(5–7), microfluidic tumor-on-a-chip devices(8–10), and scaffold-based and bio-printed models(11–15). Each model type can incorporate tumor cancer cells, tumor non-cancer cells, and extracellular matrix, thus mimicking continuous cell-extracellular matrix (ECM) and cell-cell interactions *in vivo*. Such models provide opportunities for asking specific biological questions about cancer cell interactions with their microenvironment while controlling the system in which cells are cultured. More recently, cancer patient-derived organoids (PDOs) have been introduced into 3D ECM-based models to increase cancer model fidelity by capturing important characteristics of primary tumors, such as intratumoral cancer cell heterogeneity and patient heterogeneity(16–23). 3D ECM-based cancer models are often combined with imaging-based assays, with both longitudinal and end-point readouts, as it is a non-disruptive method that enables spatial resolution, can distinguish independent cell populations, and provides granular insights into cell morphology and behavior. Furthermore, combining ECM-based cancer models with recent advances in high-content imaging and analysis provides a powerful tool to study tumor cell behavior even at a single cell level(24).

We previously reported a 3D-ECM culture platform called GLAnCE (Gels for Live Analysis of Compartmentalized Environments) that enables imaging-based assays and analysis of tumor cell dynamics, including tumor-stroma cell interactions and chemotherapy response dynamics of PDOs(25,26). In 96-GLAnCE, microliter cell-gel suspensions are molded in a 12 × 8 array of channels to generate consistent, 250 μm-thick flat microgels (Figure 1A) compatible with image acquisition using widefield high-content and confocal microscopes. The cell-gel-filled chip is then attached to a black 96-well no-bottom plate by double-sided adhesive, and complete culture media is added to each well for cell culture experiments (Figure 1A).

**Figure 1.**
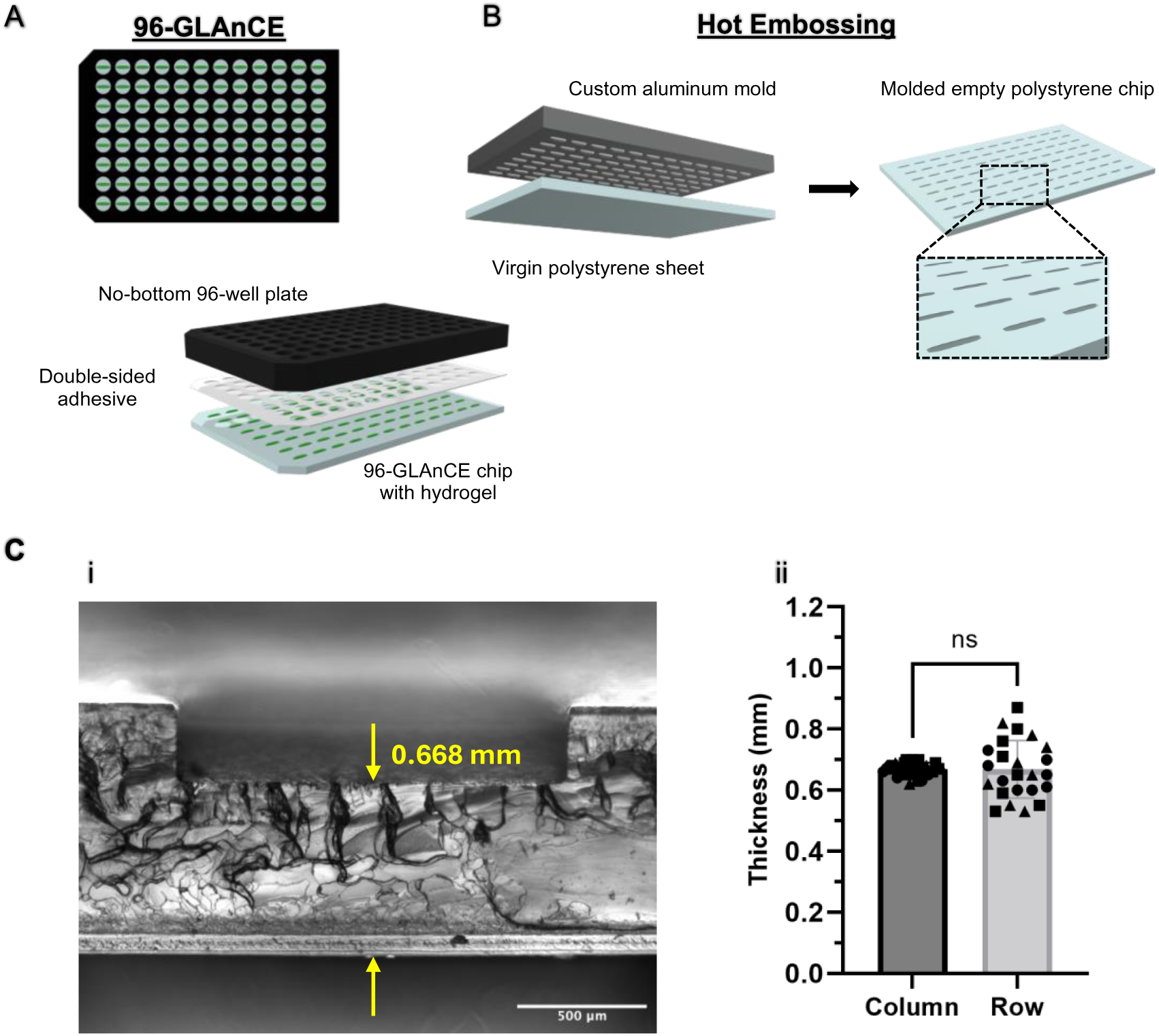
Optical limitations of the 96-GLAnCE platform manufactured by hot embossing. (A) Overview of the 96-GLAnCE platform components. A hot-embossed 96- GLAnCE chip with an array of molded cell-containing microgels is attached to a no-bottom 96-well plate using a double-sided adhesive. (B) Overview of the hot embossing fabrication technique. A virgin polystyrene sheet (1.2 mm thick) is hot-embossed using a custom positive aluminum mold with a milled 12 × 8 array of microchannels. This produces a molded, empty polystyrene chip, the 96-GLAnCE bottom component. Diagrams created with Vectary.com. (C) (i) A representative 4x brightfield image of a hot-embossed microchannel cross-section showing that after hot-embossing, the resulting chip thickness is 0.668 mm. Scale bar is 500 µm. (ii) Quantification of chip thickness within channels across all columns and rows of the platform. A caliper was used to quantify chip thickness within channels. Each plotted point represents an average chip thickness for each column/row across the platform. Statistics performed using unpaired t-test with Welch’s correction, ns – no significance. N = 3 independent 96-GLAnCE chips.

96-GLAnCE bottom chips are produced through hot-embossing, where a custom positive aluminum mold with a 12 × 8 channel array is pressed into a virgin polystyrene sheet under high temperature and pressure using a hot embossing system (Figure 1B). However, this fabrication method presents several challenges. It requires access to costly, specialized equipment and facilities, limiting the widespread adoption of the GLAnCE platform within the research community. Additionally, the method uses thick polystyrene sheets (1.2 mm) as the starting material. Our analysis revealed that the chip thickness within the channels averages 0.670 mm (±0.091), with noticeable variation across the rows (Figure 1Ci-ii). This excessive thickness falls outside the working distance range of most high-magnification objectives, restricting the use of GLAnCE devices to low-magnification lenses and preventing clear focus on samples to study cells at single-cell and subcellular levels(27).

In this study, we describe the optimization of a vacuum thermoforming method for the fabrication of 96-GLAnCE bottom chips. Thermoforming is a standard industrial process in which a plastic sheet is heated to a malleable state, then shaped over a one-sided mold using a pressure differential (28). Once cooled, the plastic is removed from the mold. While several groups have previously used pressure (micro)thermoforming to fabricate cell culture devices with microcavities from thin polymer films, these protocols still require access to hot- embossing equipment (29–31). In contrast, we demonstrate here that a commercially available desktop vacuum thermoformer—operating solely through a vacuum-generated pressure differential—can be used to produce robust 3D cell culture chips in-house, without the need for cleanroom facilities or specialized fabrication equipment. Using methods for polydimethylsiloxane (PDMS) casting and brightfield imaging, we characterize various vacuum thermoforming parameters and develop a protocol for fabricating 96-channel GLAnCE bottom chips from thin (0.192 mm) polystyrene film. We validate that these thermoformed chips are fully compatible with the standard 96-GLAnCE platform assembly and seeding workflows. Furthermore, we demonstrate that the thermoformed GLAnCE devices support cancer cell proliferation with minimal gel contraction and are compatible with three different gel matrices. To showcase the further capabilities of these thermoformed chips, we culture primary pancreatic fibroblasts and pancreatic cancer patient-derived organoids (PDOs), perform *in situ* immunofluorescent staining, and acquire images using confocal microscopy and high-magnification objectives (up to 63x) to visualize cell features at single-cell and subcellular levels. These results highlight the potential of vacuum thermoforming as a flexible, cost-effective method for manufacturing 3D cell culture platforms for enhanced imaging applications.

## 2. Materials and methods

### 2.1 Cell culture

The KP4 pancreatic cancer cell line (JCRB Cellbank) was maintained in IMDM supplemented 10% v/v fetal bovine serum (FBS) and 1% v/v penicillin and streptomycin (Wisent Inc., Canada) and passaged every three days. Pancreatic stellate cells (PSCs) (reference #3830, ScienceCell) were maintained in DMEM containing 10% v/v FBS (Wisent, Canada) and 1% v/v penicillin and streptomycin (Wisent, Canada). Medium was changed every three days, and cells were passaged every 6–7 days. PSCs were used for a maximum of 8 passages. Pancreatic ductal adenocarcinoma organoids established from PDAC patients were obtained from the UHN Biobank at Princess Margaret Cancer Center (PMLB identifier PPTO.46 from the University Health Network, Ontario, Canada) in compliance with the University of Toronto Research Ethics Board guidelines (protocol #36107). PPTO.46 organoid cultures were maintained in Growth Factor Reduced Phenol Red-Free Matrigel™ Matrix domes (Corning Life Sciences, Corning, USA) with 500 μL of complete growth medium as previously reported (12,26) . PPTO.46 organoid cultures were passaged weekly (1:4 split ratio) and used for a maximum of 30 passages. All cells were tested for mycoplasma every three months. All cultures were maintained in a humidified atmosphere at 37 °C and 5% CO2. All cells were tested for mycoplasma every three months.

### 2.2 Lentivirus production and cell transduction

Green fluorescence protein (GFP) lentivirus was produced using calcium phosphate co-transfection of HEK293T cells as previously described(12,32). Wild-type KP4 cells and wild-type PSCs were transduced with concentrated virus, and GFP-expressing cells were selected using puromycin (1 μg/mL and 1.5 μg/mL for KP4 and PSCs, respectively, Sigma-Aldrich, St Louis, USA).

### 2.3 Fabrication of custom device components

Positive aluminum molds were fabricated by micro-milling an array of 3 × 3 or 12 × 8 channel-shaped features (6 mm × 1.5 mm × 250 μm) into the surface of aluminum slabs (6061, T6, McMaster-Carr). The features were polished using fine-grit sandpaper to remove any imprints. Finally, vacuum holes were drilled around the channels. Vacuum hole configurations for 3 × 3 (configurations 1-9) and 12 × 8 molds are specified in **Table 1**.

**Table 1.** Summary of vacuum hole configurations.

| Vacuum hole configuration | Number of vacuum holes per channel | Vacuum hole distance to channel (mm) | Vacuum hole diameter (mm) |
| --- | --- | --- | --- |
| Channel 1 | 2 | 1.0 | 1.0 |
| Channel 2 | 4 | 1.0 | 1.0 |
| Channel 3 | 6 | 1.0 (0.2 for side holes) | 1.0 |
| Channel 4 | 2 | 0.2 | 1.0 |
| Channel 5 | 4 | 0.2 | 1.0 |
| Channel 6 | 6 | 0.2 (0.2 for side holes) | 1.0 |
| Channel 7 | 2 | 0.2 | 0.5 |
| Channel 8 | 4 | 0.2 | 0.5 |
| Channel 9 | 6 | 0.2 (0.2 for side holes) | 0.5 |
| **12 × 8 mold** | **4** | **0.2** | **0.5** |

PDMS slabs, used for cell seeding, and double-sided adhesive, used to attach cell-filled GLAnCE chips to the black no-bottom 96-well plate (Greiner Bio-One), were prepared as previously described(25,26). In brief, SYLGARD 184 Silicone Elastomer base and curing agent were mixed in the recommended 9:1 ratio (Sylgard 184, Ellsworth Adhesives), degassed for 30 min in a vacuum, and cured in rectangular plastic molds at 60 °C overnight. The 96 inlet and 96 outlet holes were aligned in PDMS slabs and punched out using a 1 mm bio-punch (Integra). Double-sided adhesives were prepared by cutting non-toxic adhesive tape (ARcare, Adhesive Research) to the dimensions of a standard 96-well plate using a Silhouette Cameo Electronic Cutting Machine (Silhouette).

PDMS was additionally used to fabricate structural support of the thermoformed 96-GLAnCE during device assembly. A thermoformed 96-GLAnCE chip was placed face-down at the bottom of a 9cm × 13cm plastic mold. 50g of PDMS was prepared using the abovementioned methods, poured over the PS chip in the plastic mold, and cured at 60°C overnight. After curing, the PS chip was detached from the PDMS support, revealing an array of channels imprinted into the PDMS slab.

### 2.4 Vacuum thermoforming of polystyrene film and GLAnCE chips

The vacuum thermoforming process is depicted in Figure 2A; the desktop thermoformer (Supplementary Figure 1A) was first preheated at the temperature setting (specified in Table 2) for 10 minutes. The positive aluminum molds were always kept in the oven at 60 °C and preheated to the temperatures (specified in Table 2) for 1 hour. A 250 mm-by-350 mm sheet of 0.192 mm-thick PS film (Figure 2A) was cut from a roll of PS film (Sigma Aldrich, USA). After 10 minutes, the sheet of PS film was clamped and lifted to the heating element (Figure 2Aii) to be heated for the selected timings (specified in Table 2). 15 seconds before the PS film heating was completed, the vacuum was turned on, and the mold was taken out of the oven and placed in the center of the thermoformer vacuum bed. Once the PS film heating was completed, the clamped PS film was lowered over the mold to activate the vacuum (Figure 2A). After the vacuum was turned off, the thermoformed PS film was left to cool down for 5 min, unclamped, and retrieved from the mold. For subsequent PDMS casting, thermoformed film samples were cut out, leaving a 1-cm border, and placed inside plastic dishes. Cut-out thermoformed chips were trimmed using a scalpel to 115 mm × 80 mm dimensions for seeding experiments and exposed to UV light for 2.5 h before use.

**Figure 2.**
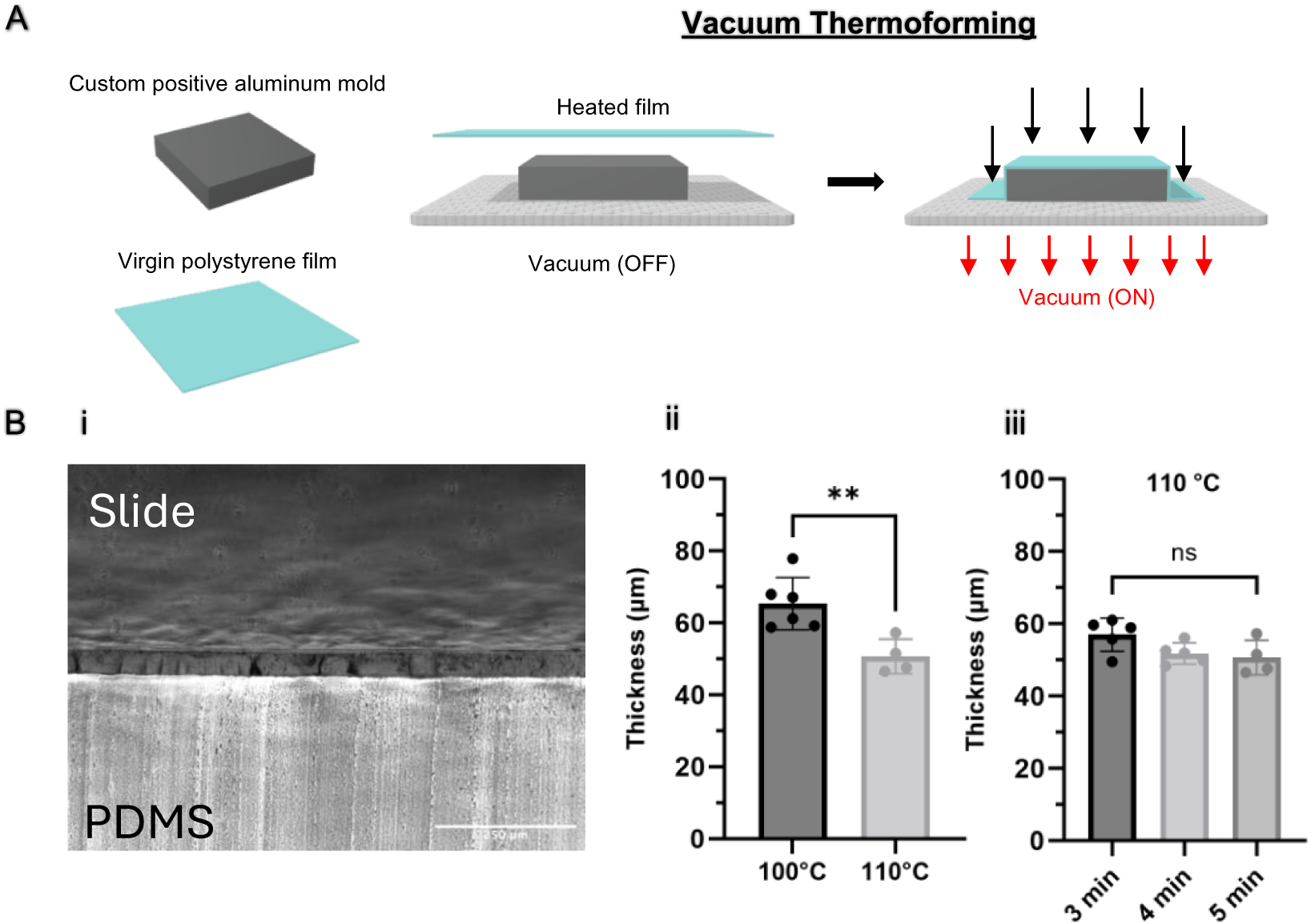
Selecting parameters for vacuum thermoforming of polystyrene film. (A) Overview of vacuum thermoforming working principle. A custom positive aluminum mold is placed onto a vacuum bed. A polystyrene film is heated to a temperature within its thermoforming window or rubbery plateau region and then drawn over the mold. This activates the vacuum that pulls the malleable film tightly against the mold surface. When the thermoformed film cools down and is retrieved from the mold, it preserves the shapes and patterns of the mold. Diagrams created with Vectary.com. (B) (i) A representative 10x brightfield image of a polystyrene film cross-section used to quantify film thickness after thermoforming. Scale bar is 250 µm. (ii) Quantification of film thickness after heating at 100 and 110 °C temperature settings for 5 minutes. Each plotted point represents an independent film sample. Statistics performed using unpaired t-test using Welch’s correction, **p< 0.01. N= 4-6 independent film samples. (iii) Quantification of film thickness after heating at 110 ° C for 3, 4, or 5 minutes. Each plotted point represents an independent film sample. Statistics performed using ordinary one-way ANOVA, ns – no significance. N= 4-5 independent film samples.

**Table 2.** Summary of tested vacuum thermoforming parameters.

| Type of characterization | Thermoformer heating setting | PS film heating time | Mold pre-heating temperature |
| --- | --- | --- | --- |
| Thermoformed PS film | 100 °C | 3 min | 60 °C |
| (3 × 3 mold, flat side) | 110 °C | 4 min |  |
|  |  | 5 min |  |
| Thermoformed chip with channels | 110 °C | 4 min | 120 °C |
| (3 × 3 mold, side with channels) |  |  | 150 °C |
| Thermoformed 96-GLAnCE chip | 110 °C | 4 min | 150 °C |
| (12 × 8 mold) |  |  |  |

### 2.5 Characterization of polystyrene film thickness and channel dimensions using PDMS casting and brightfield imaging

An overview of the PDMS casting and brightfield imaging methods for the characterization of thermoformed PS film thickness and channel dimensions is depicted in Supplementary Figure 1C. PDMS solution was prepared using SYLGARD 184 Silicone Elastomer Kit according to the manufacturer’s recommended base-to-curing agent ratio (9:1) (Sylgard 184, Ellsworth Adhesives). The mixture was degassed for 30 min in a vacuum, then poured into prepared thermoformed samples in plastic dishes and cured at 60 °C overnight. 4 and 30g of the PDMS solution were required to fully cover one sample of thermoformed film/chip with channels (3 × 3) and 96-GLAnCE chip (12 × 8 mold), respectively.

After curing overnight, thermoformed samples with casted PDMS were cut with a razor blade into smaller flat film samples or individual channels. Each film sample/channel was bisected to obtain two duplicate cross-sections. One duplicate cross-section of each sample was imaged using a BF microscope with MetaMorph Premier (v7.7.5.0), using an exposure time of 100 ms. The microscope images were then analyzed using straight line (for film thickness and channel depth) and angle (for channel side wall angle) selection tools in FIJI (v1.54f), using the appropriate pixel-to-distance scale for the magnification used.

To measure PS film thickness, samples were imaged at 10x magnification using an Olympus IX81 microscope. The scale was set at 1.52 pixel/μm. The straight-line tool was used to measure the thickness of the PS film at three random locations on the image to compute an average sample thickness. To measure channel wall angle and depth, the channel samples were imaged at 4x magnification using an Olympus IX81 microscope, and using the scan slide functionality of MetaMorph Premier, to obtain three images containing the channel and the beginning of the flat region of the PS film on either side of the channel. The scale was set at 0.624 pixel/μm. The three images were stitched with an automated macro using the pairwise stitching plugin in FIJI with the linear blending method(33). A reference line was drawn in FIJI between the flat regions on either side of the channel. The angle tool was then used to obtain the angle of both channel walls relative to the reference line, and the two angles were averaged to obtain the average channel wall angle of the sample. The channel depth was determined by measuring the distance between the reference line and the channel bottom using the straight-line tool at three random locations along the channel and computing an average.

### 2.6 Inter-well leakage assessment

Methods for the inter-well leakage assessment were adapted from the previously published work(12). The thermoformed 96-GLAnCE chip was aligned and securely attached to a black no-bottom 96-well plate with a double-sided adhesive; 200 μL of 1% v/v fluorescein solution (in PBS) or blank control (PBS) was added to alternating wells in a checkerboard pattern (Figure 3). The fluorescence of each well was measured using a microplate reader (Tecan Pro) immediately after assembly and again after 24 hours of incubation in the dark.

**Figure 3.**
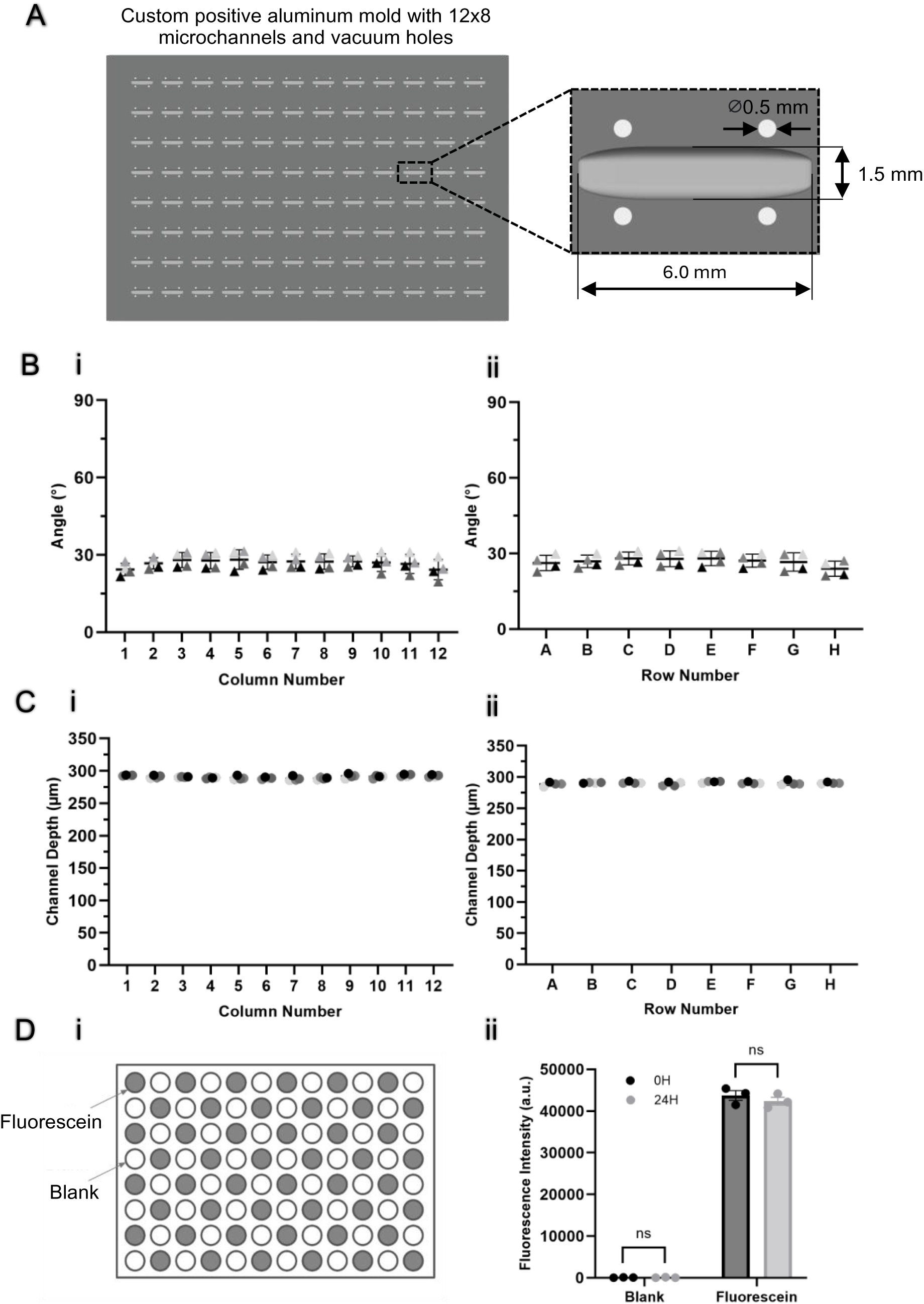
Characterization of thermoformed 96-GLAnCE chips. (A) Design of a custom positive aluminum mold with a 12 × 8 array of microchannels and vacuum holes from Channel 8 vacuum hole configuration. Diagrams created with Vectary.com. (B) Quantification of channel angle of thermoformed 96-GLAnCE chips across (i) columns (1-12) and (ii) rows (A- H). Each plotted point represents an independent chip. Statistics performed using ordinary one- way ANOVA. N = 4 independent thermoformed 96-GLAnCE chips. (C) Quantification of channel depth of thermoformed 96-GLAnCE chips across (i) columns (1-12) and (ii) rows (A- H). Each plotted point represents an independent chip. Statistics performed using ordinary one- way ANOVA. N = 4 independent thermoformed 96-GLAnCE chips (D) (i) Checkerboard layout for the interwell leakage test. (ii) Quantification of fluorescence levels in blank (wells with PBS) and fluorescein wells of the thermoformed 96-GLAnCE platform at 0h and 24h. Each plotted point represents an average of 48 wells per condition. Statistics performed using paired t-test, ns – no significance. N = 3 independent thermoformed 96-GLAnCE chips.

### 2.7 Thermoformed 96-GLAnCE seeding and assembly

The thermoformed 96-GLAnCE seeding and device assembly protocol were adapted from the original workflow with minor changes (Figure 4)(26). A PDMS slab with bio-punched holes was aligned with channels and firmly attached to the thermoformed chip. The PDMS slab-chip construct was then placed on the PDMS support, and the chip channels were aligned with the imprinted channels on the PDMS support. This construct was kept at 37 °C until use. 3 mg/mL of rat tail collagen type I was prepared according to the manufacturer’s protocol (Ibidi, USA) and, after neutralization, kept on ice and used within 15 minutes. Fluorescent FITC beads (Bangs Laboratories, FCDG009) were resuspended in the gel to yield a concentration of 10,000 beads/6.5 μL plus an excess of 50% to account for higher gel viscosity and small pipetting volumes(25). Varying volumes (for gel volume testing experiments) or 3.6 μL (for full-chip seeding) of bead-gel suspension were injected through the inlet of each closed channel of the GLAnCE seeding device. Once seeding was complete, the seeding device with gel-filled channels was kept at 37 °C for 30 min. For the GLAnCE platform assembly, the PDMS support aided the detachment of the PS chip from the PDMS slab, after which the PS chip was removed from the PDMS support. Finally, the PS chip was aligned and securely attached to a black no-bottom 96-well plate with a double-sided adhesive, and 200 μL of PBS was added to each well. A standard 96-well plate lid (Greiner Bio-One) was used as the GLAnCE lid; the assembled GLAnCE platform was finally placed over another standard 96-well plate to prevent damage to the thin thermoformed PS chip.

**Figure 4.**
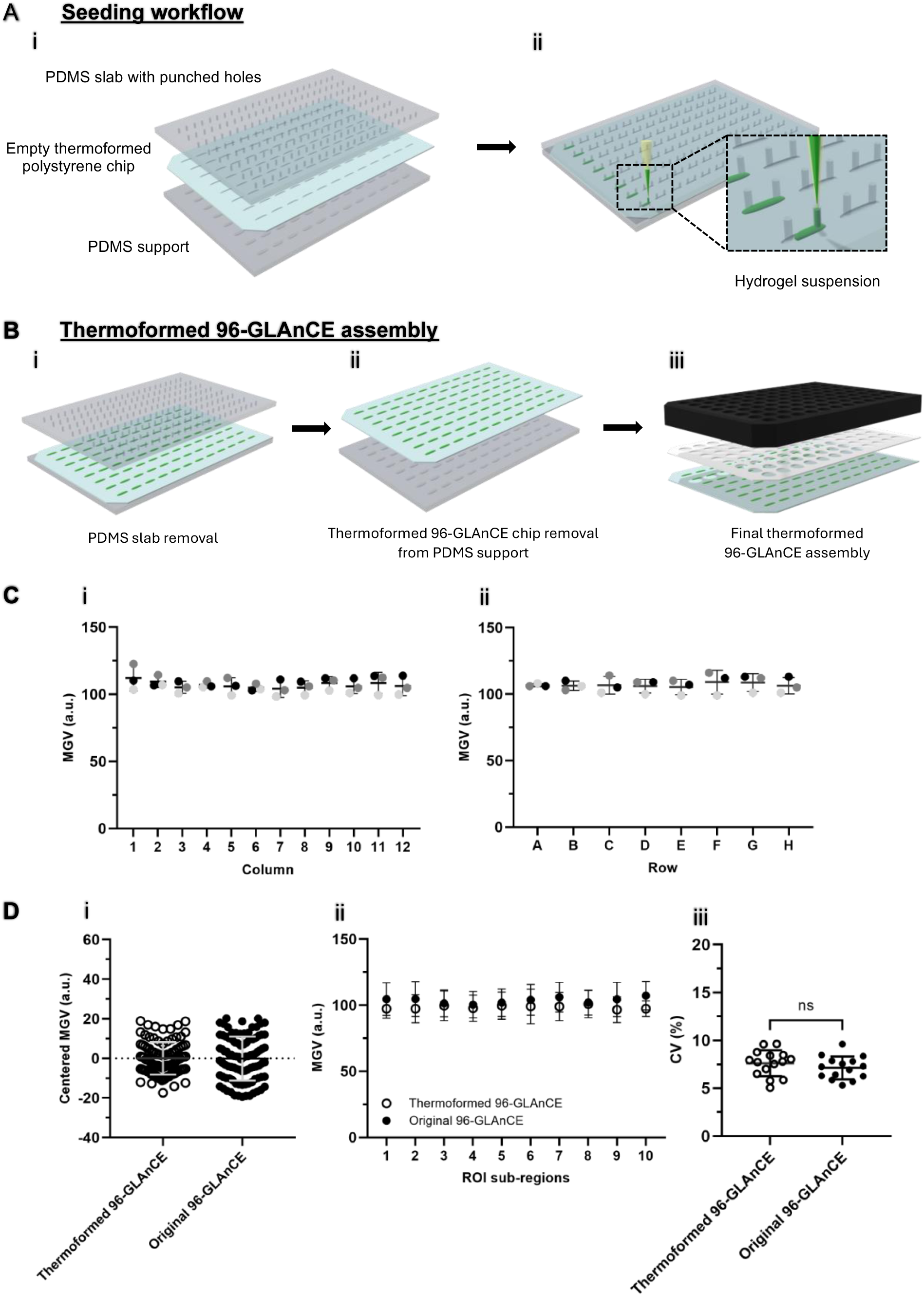
Gel seeding characterization in thermoformed 96-GLAnCE chips. (A) Overview of gel seeding workflow adapted for thermoformed 96-GLAnCE chips. (i) The seeding device consists of a thermoformed chip aligned with a PDMS slab with inlet and outlet holes on the top and a PDMS support with molded 96 channels on the bottom. (ii) Hydrogel suspension is injected through an inlet hole into the channel for all 96 channels and left to gel at 37 °C. (B) Overview of the final device assembly workflow. (i) After gelation, PDMS slab is horizontally detached from the thermoformed chip with microgels. (ii) Next, the thermoformed chip with microgels is horizontally detached from the PDMS support and (iii) immediately aligned and attached to a 96-well black no-bottom plate using a double-sided adhesive. Diagrams created with Vectary.com. (C) Quantification of mean gray value (MGV) in thermoformed 96- GLAnCE chips across (i) columns (1-12) and (ii) rows (A-H). Each plotted point represents an independent seeding experiment. Statistics performed using ordinary one-way ANOVA. N = 3 independent seedings. (D) (i) Interwell variation for full-seeded thermoformed and original 96- GLAnCE chips. Error bars are SD for 96 technical replicates; MGV was centered to the mean for each chip type. (ii) Quantification of spatial homogeneity of gel spread and the spatial spread across ROIs for each chip type. Each ROI was divided into 10 even sub-regions from left to right, and the MGV was measured for each sub-region. Results show mean ± SD for 15 channels (technical replicates). (iii) Quantification of the coefficient of variation between the 10 sub-regions within one ROI to characterize intrawell seeding variation for each chip type. Results show mean ± SD for 15 channels (technical replicates). Statistics performed using unpaired t-test with Welch’s correction, ns – no significance.

### 2.8 Quantification of gel profiles and seeding reproducibility in original and thermoformed GLAnCE using an image-based readout

Microgels in the assembled thermoformed GLAnCE were imaged using a high-throughput widefield microscope IX Pico (Molecular Devices, USA) with the following settings (4x objective; FITC channel exposure time = 10 ms; autofocus was set on well bottom using a wide search range). Image analysis and quantification were performed in FIJI (v1.54f). The scale was set at 0.580 pixel/μm

*Gel volume testing:* Channels were seeded with 3.0, 3.3, or 3.6 μL of FITC bead-gel suspension, and widefield images (2 sites) were acquired to capture the whole channel and automatically stitched using IXP software. The images were then thresholded using the FIJI default algorithm, and the ROI of channel dimensions was applied to quantify the bead area coverage relative to the ROI area.

*Inter-well heterogeneity comparison for full-chip seeding:* thermoformed and original GLAnCE chips were seeded with 3.6 and 3.0 μL of FITC bead-gel suspension, respectively. Widefield images of the central part of channels (1 site) were acquired to remove GLAnCE seeding artifacts (Supplementary Figure 5B) and reduce image acquisition time. A centrally positioned 1100 × 700 ROI was applied to each 96 channels to quantify the mean gray value (MGV).

*Intra-well heterogeneity comparison:* for both thermoformed and original GLAnCE chips, the 1100 × 700 ROI, centrally positioned in a channel, was cropped and divided into 10 even sub- regions from left to right for 15 randomly selected channels. The MGV of each sub-region was measured to compute the standard deviation within the cropped ROI; the standard deviation was then divided by the MGV of each sub-region to obtain the coefficient of variation associated with the cropped ROI, a metric previously used by our group to assess seeding homogeneity within the channel(26).

### 2.9 Quantification of cell line growth in different seeding hydrogel matrices in thermoformed GLAnCE

We assessed the thermoformed GLAnCE chips compatibility for longitudinal cell proliferation assay in three different gel matrices: Matrigel (Growth Factor Reduced Phenol Red-Free Matrigel™ Matrix (Corning Life Sciences, Corning, USA)), Bovine Collagen Type I (3mg/mL; PureCol, Advanced BioMatrix), and Rat Tail Collagen Type I (3mg/mL, Ibidi). Rat Tail Collagen Type I solution was prepared as described above. Bovine Collagen Type I solution was prepared as previously reported(12). In short, bovine collagen hydrogel was prepared by mixing 8 parts 3 mg mL^−1^ type I bovine collagen with 1 part 10× minimal essential medium (MEM, Life Technologies, Grand Island, USA) by volume and neutralizing to pH 7 endpoint with 0.8 M NaHCO3 (Sigma-Aldrich). The hydrogel solutions were kept on ice until the time of use. The adherent GFP^+^ KP4 cells were trypsinized following a standard protocol and pelleted by centrifugation (300 g, 5 min, 4 °C). 1500 cells μL^-1^ of GFP^+^ KP4 cells were resuspended in the required volumes of prepared gel solutions plus an excess of 50% over ice and then seeded into the thermoformed GLAnCE chip as described above. Since all three gels have different gelation times, GFP^+^ KP4 cells were seeded into separate thermoformed GLAnCE chips for each gel type. Matrigel, Rat Tail Collagen Type I, and Bovine Collagen Type I were incubated at 37 °C for 10, 30, and 45 minutes, respectively. Complete cell culture medium was added to each well, and all conditions were imaged using a high-throughput widefield microscope (4x objective; FITC channel exposure time = 10 ms; autofocus was set on the well bottom using a wide search range) immediately after the seeding and on days 2, 4, and 6. Note that we observed some variation in the intensity of the GFP signal of KP4 cells between independent seedings. To account for this variation, we compared fold changes in MGV instead of raw values for each condition.

### 2.10 Immunofluorescence staining and confocal imaging

We assessed the thermoformed GLAnCE chips’ suitability for confocal microscopy with high-magnification objectives by imaging GFP^+^ KP4 cells and performing immunostaining and imaging of patient-derived organoid PPTO.46 and GFP^+^ PSCs. All three cell types were seeded in Rat Tail Collagen Type I (3mg/mL). For GFP^+^ KP4 cells, cells were seeded at the density of 3000 cells μL^-1^ and cultured for 3 days. For PPTO.46, cells were seeded at the density of 1500 cells μL^-1^ and cultured for 5 days. Before seeding, PPTO.46 dissociated cells were strained through 70 μm cell strainers (Fisherbrand) to remove large clumps. For GFP^+^ PSCs, cells were seeded at the density of 6000 cells μL^-1^of Rat Tail collagen and cultured for 1 or 4 days. All cells were fixed inside GLAnCE channels with 4% PFA for 10 minutes at room temperature (RT). Cytokeratin 19 (CK19) immunostaining of PPTO.46 cells was done as previously described(26). Briefly, cells were permeabilized with 0.5% Triton-X 100 (Millipore Sigma) for 5 min, blocked with 5% BSA in IF buffer (0.1% BSA/0.2% Triton-X 100/0.05% Tween-20) for 30 min at RT, and then incubated with CK19 (Abcam, Catalog# ab9221) primary antibody (Abcam, Catalog# ab9221) in the blocking buffer at 4 °C overnight. The secondary antibody conjugated to Alexa Fluor 594 (Thermo Fisher Scientific) was incubated for 2 hours, followed by the nuclear counterstaining with DRAQ5 (Cell Signalling Technology) for 10 min at RT . For staining of cytoskeletal fibers and intracellular lipid droplets of GFP^+^ PSCs, microgels were permeabilized with 0.1% Triton X-100 for 5 min and then blocked with 2.5% BSA for 30 min. For cytoskeleton staining, Alexa Fluor 647 conjugated α-SMA antibody (ab196919, Abcam) was diluted 1:200 and incubated overnight at 4 °C. For lipid droplet staining, microgels were incubated with 1 μg ml^−1^ BODIPY Red (Invitrogen) solution in PBS for 30 min at room temperature. All images were acquired using Leica SP8 confocal, equipped with the following objectives: 10x objective (NA 0.4), 20x oil objective (NA 0.75), and 63x oil objective (NA 1.40).

### 2.11 Statistics

Statistical analysis was carried out in GraphPad Prism 10 (GraphPad Software, USA). An unpaired student t-test with Welch’s correction was used to evaluate the original 96-GLAnCE channel thickness, thermoformer heating temperatures, mold pre-heating temperatures; channel depth, autofluorescence, and intra-well seeding variation in original and thermoformed 96-GLAnCE chips. A paired t-test was used to assess inter-well leakage. One-way ANOVA with Tukey’s HSD were applied to assess thermoformer heating timings; channel wall angle and depth characterization in thermoformed 96-GLAnCE chips; gel volume testing and inter-well seeding variation in thermoformed GLAnCE chips; and GFP^+^ KP4 cell proliferation in three different matrices. *p* < 0.05 was considered significant.

## 3. Results and discussion

### 3.1 Optimization of vacuum thermoforming heating regime for polystyrene films

In this study, we set out to develop methods to rapidly manufacture 96-GLAnCE bottom chips using vacuum thermoforming. Vacuum thermoforming is a standard industrial process for shaping plastics, in which a sheet of plastic is heated until malleable, and a vacuum-based pressure differential is used to tightly fit the malleable plastic over a one-sided mold (Figure 2A)(28,34). When the thermoformed plastic cools down and is retrieved from the mold, it preserves the shapes and topographic patterns of the mold. This study used a portable desktop Mayuku Formbox thermoformer (Supplementary Figure 1A) connected to a shop vacuum. The desktop thermoformer comprises a heating element, a vacuum bed, and trays. The vacuum bed holds a custom mold, allowing vacuum suction across the mold surface. The trays enable a user to secure a plastic material and bring it up close to the heating element, which heats it to a malleable state.

Film heating temperature is a critical parameter in vacuum thermoforming, as the plastic material needs to be at a temperature within its thermoforming window or rubbery plateau region, which occurs in a range of temperatures above the glass transition temperature where the elastic modulus does not significantly change with temperature(28). Therefore, we first sought to identify an appropriate heating regime (heating temperature and time(35)) to enable thermoforming of polystyrene. Polystyrene (PS) is used for our original hot-embossed GLAnCE chips and has a reported glass transition temperature of 100 °C (36). Because vacuum thermoforming is compatible with much thinner plastic films as a starting material, we opt for the 0.192 mm-thick polystyrene film instead of much thicker starting sheets used in our original device (1.2 mm). We first tested the two thermoformer temperature settings at and above the glass transition temperature of polystyrene (settings 5 and 6 corresponding to 100 and 110 °C, respectively (Supplementary Figure 1B) and characterized the resulting thickness of thermoformed films by using methods for PDMS casting and brightfield imaging (Supplementary Figure 1C). Film samples heated to 110 °C for 5 mins were significantly thinner than films heated at 100 °C (50.7 μm±4.9 vs 65.4 μm±7.3; Figure 2Bii), consistent with the higher temperature being above the material glass transition temperature. When using 110°C, reducing the PS film heating time from 5 to 4 and 3 minutes did not significantly change the thermoformed film thickness; however, we observed the thickness was more consistent when using a heating time of 4 minutes (51.8 μm±2.9; Figure 2Biii). Therefore, we selected a heating temperature and time of 110 °C and 4 minutes as standard operating conditions for further tests.

For the scope of this study, we only attempted thermoforming of PS films to keep the material consistent with the original 96-GLAnCE platform, in which PS was selected for its ability to bind non-specifically to natural polymers, thus improving immobilization of the microgels within the chip channels. Our fabrication method could be re-optimized for other transparent thermoplastic polymers depending on the specific device application. However, when using a different polymer film, the main limiting factor to consider is the heating temperature range of a desktop thermoformer, which must accommodate temperatures around the polymer’s glass transition temperature for successful manufacturing. For example, our specific desktop thermoformer has an upper-temperature setting of 110 °C, which is not high enough for thermoforming polycarbonate films with a glass transition temperature of around 140 °C (37), but can work for acrylic polymers and cyclic olefin copolymers with lower norbornene content (for these materials, glass transition temperatures typically range from 80-105 °C) (38,39).

### 3.2 Optimization of the mold design and heating parameters using a 3 × 3 prototype

Having identified heating parameters for sufficient PS film thermoforming, we next focused on the positive mold design and mold heating regime. We hypothesized that choosing an appropriate mold preheating temperature and incorporating vacuum holes around channels (number and position) would produce well-defined channels for molding microgels of consistent thickness across the chips. We therefore designed a custom positive aluminum mold with a 3 × 3 array of channels with nine vacuum hole configurations (labeled from 1 to 9) (Supplementary Figure 2A). We tested each vacuum hole configuration at two mold preheating temperatures (120 °C and 150 °C). The specific dimensions of each vacuum hole configuration are provided in Table 1. For each configuration design and preheating temperature, we measured the resulting channel wall angle using the flat region on either side of the channel cross-section as a reference line (Supplementary Figure 2Bi). Molds preheated to 120 °C resulted in channels with poorly defined profiles for all vacuum hole configurations with an average channel wall angle of 11.7°±0.3 (Supplementary Figure 2Bii). Increasing the mold preheating temperature to 150°C progressively increased the mean channel wall angle to 30.6°±1.6. However, any further increase in the mold preheating temperature led to the sample warping (data not shown). Therefore, we continued using 150°C as a standard operating temperature for mold preheating.

For vacuum hole configurations, designs containing 1 mm-diameter vacuum holes (Supplementary Figure 2A, #1-6) produced visible indentations in the thermoformed films. These indentations prevented the complete sealing of thermoformed channels by a PDMS slab, thus rendering gel seeding impossible in the preliminary seeding experiments (data not shown). However, the designs with 0.5 mm-diameter vacuum holes (Supplementary Figure 2A, #7-8) produced only minor indentations that enabled effective channel sealing for gel seeding. Of the designs with 0.5 mm holes, configuration #8 consistently generated channels with the greatest wall angle (32.0°±2.5); therefore, we selected this configuration to produce a scaled-up positive mold.

### 3.3 Scale-up of thermoforming mold to enable 96-GLAnCE chip manufacturing

Using vacuum hole configuration #8, we generated a positive mold with a 12 × 8 array of channels with four 0.5 mm vacuum holes drilled around each channel for thermoforming 96- GLAnCE chips (Figure 3A). Thermoformed channels showed consistent wall angle in a single thermoformed 96-GLAnCE chip and across multiple chips (Figure 3Bi-ii), with a slight decrease observed at the chip’s periphery (24.3°±3.0 vs 27.3.2°±2.6). This decrease is likely because the periphery channels are located farther from the vacuum source; therefore, the deformation pressure to adhere the polymer sheet to the mold is lower. To determine if this decreased wall angle affected other channel dimensions, we also characterized the channel depth of thermoformed chips across columns and rows (Figure 3Ci-ii). We did not observe a decrease in the channel depth at the chip’s edge; the depth appeared consistent across the 96 channels in a single thermoformed chip and across multiple chips (across columns - 290.4 μm±1.75; across rows - 290.3±1.2). Notably, the channel depths in thermoformed chips were comparable to the ones in original hot-embossed 96-GLAnCE chips (across columns 290.9 μm±4.9; across rows – 291.7 μm±4.0; Supplementary Figure 3).

In addition to characterizing the variation in channel dimensions, we confirmed the thermoformed 96-GLAnCE chips were compatible with the original device assembly workflow by assessing inter-well leakage. To do this, we assembled 96-GLAnCE plates using thermoformed chips, added fluorescein solution to alternating wells in a checkerboard pattern (Figure 3Di), and measured fluorescence immediately after the device assembly (0 hours) and after 24 hours. No fluorescein leakage was observed between the wells at the 24-hour timepoint, suggesting the thermoformed chips could be robustly assembled into 96-GLAnCE plates with individually addressable culture wells.

Our work showcases that vacuum thermoforming using a commercially available desktop thermoformer offers a highly accessible alternative to hot embossing for in-house fabrication of cell culture devices of mm-scale dimensions. Further, although we only focused on forming mm-scale straight channels for this study, we anticipate more advanced geometric shapes of the same scale would be possible with novel custom aluminum molds. Furthermore, since the thermoforming process does not involve material melting phase, it is compatible with polymers with surface or bulk physicochemical modifications(30,40,41). Vacuum thermoforming could generate scaffolds with tissue-like structures, topographical cues, and 3D confinements for applications beyond tissue modeling, including tissue development and mechanobiology studies(42,43). Note that the fabrication of devices with features of micro-scale resolution will likely require more advanced approaches, such as pressure (micro)thermoforming using hot- embossing equipment or other micro moulding and lithographic methods(29,31,44,45).

### 3.4 Validation of thermoformed chips in the current 96-GLAnCE platform seeding and imaging workflows

The original hot-embossed 96-GLAnCE platform has been previously applied for imaging-based readouts using widefield fluorescence microscopy. Therefore, before applying the new thermoformed 96-GLAnCE platform for further imaging workflows, we assessed whether thermoformed chips were comparable to the original ones in autofluorescence in conventional fluorescence channels and thus compatible with high-quality real-time imaging. When exposed for 300 ms, the thermoformed chips showed lower autofluorescence across all channels, especially the DAPI channel, where the autofluorescence was reduced nearly twofold (Supplementary Figure 4A,B).

Another important characteristic of the original 96-GLAnCE platform is that it requires a low seeding gel volume, significantly reducing experimental costs(26). Given this, we determined the minimum gel volume needed to maintain consistent seeding across channels in thermoformed chips. Since the channel dimensions slightly increased due to the non-vertical channel wall angle, we anticipated that the gel volume necessary for consistent channel filling would be greater than the previously used 3.0 µL. We, therefore, tested several volumes of FITC bead-gel suspension (3.0 µL; 3.3 µL (10% increase); and 3.6 µL (20% increase)) (Supplementary Figure 5Ai-ii). As expected, 3.3 and 3.6 µL exhibited greater channel area coverage when compared to 3.0 µL. However, only 3.6 µL gel volume produced consistent channel coverage across wells in a single chip and independent seedings (Supplementary Figure 5Aii). For 3.6 µL seeding gel volume, we also observed the previously reported seeding artifacts, such as inlet holes and outlet ‘tails’, caused by excess gel filling holes of the removable PDMS slab (26) (Supplementary Figure 5Ai). This further indicated that using a 3.6 µL seeding volume best mimicked the original 96-GLAnCE. Note that to prevent these artifacts from affecting further image analysis, we defined a region of interest (ROI) (Supplementary Figure 5B, ROI highlighted by a yellow dashed rectangle) consistently for all microgels for analysis, as for the original 96-GLAnCE (26). The channel ROIs for thermoformed and original 96-GLAnCE platforms appeared visually comparable (Supplementary Figure 5Bi-ii).

Having selected an appropriate gel volume and an ROI, we next seeded an entire thermoformed 96-GLAnCE chip with FITC bead-gel suspension to benchmark the seeding variation against the original 96-GLAnCE. The full-plate seeding was essential to confirm that the thermoformed 96-GLAnCE is suitable for medium-throughput assays requiring low inter-well and user-induced variation. However, we had to adapt the previously reported seeding workflow and introduce a PDMS support below a much thinner thermoformed chip to provide structural support during seeding (Figure 4Ai) and aid the removal of the PDMS slab from the thermoformed chip during the final device assembly (Figure 4Bi-ii). To quantify seeding variation across wells in a single thermoformed chip and across multiple chips, we selected the mean gray value (MGV) as an image-based metric for our ROIs since it captures signal intensity across the z-planes in widefield microscopy (26). We detected no significant differences in the inter-well seeding variation across columns and rows of the thermoformed chip for three independent seedings (Figure 4Ci-ii), suggesting our selected seeding parameters and workflow were sufficiently robust. We also benchmarked this inter-well variation against the original 96-GLAnCE by comparing all 96 ROIs and detected no substantial differences in the levels of well-to-well variation (Figure 4Di). Lastly, using the same MGV metric, we evaluated the intra-well seeding variation in thermoformed and original 96-GLAnCE chips. We divided ROIs from 15 randomly selected channels into 10 even subregions from left to right and then obtained and plotted MGV for each sub-region (Figure 5Dii). Both chip types showed flat, homogeneous bead-gel profiles. The homogeneous bead distribution was further confirmed by comparable mean coefficients of variation (CV) between the 10 sub-regions (7.6% vs. 7.1% for thermoformed and original chips, respectively) (Figure 4Diii). Together, this data suggested that thermoformed chips were compatible with the seeding workflow of the previously validated hot-embossed chips and could be applied for further imaging-based cell assays in the 96-GLAnCE platform.

**Figure 5.**
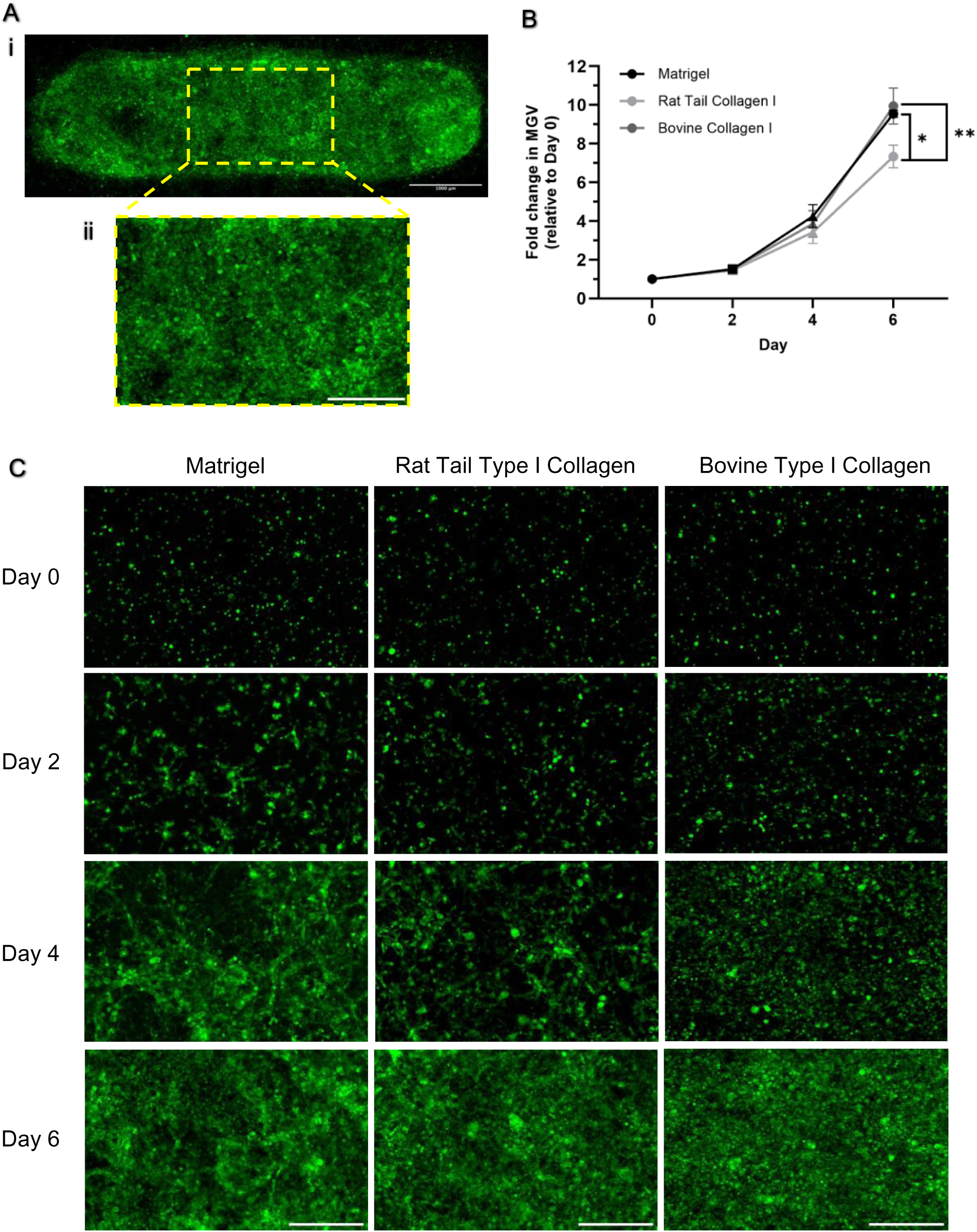
Characterization of cell proliferation in different matrices in thermoformed 96- GLAnCE. (A) (i) Representative widefield image (2 sites) of GFP^+^-KP4 cells on Day 6 after seeding in a thermoformed 96-GLAnCE chip and (ii) its cropped region of interest (ROI, 1100 × 700 pixels) used to quantify cell proliferation. Scale bars are 1000 and 500 µm, respectively. (B) Quantification of GFP^+^-KP4 cell proliferation using mean gray value (MGV) in Matrigel, Rat Tail Collagen Type I, and Bovine Collagen Type I over 1 week. Statistics performed using ordinary one-way ANOVA;*, p< 0.05; **, p< 0.01. N = 3 independent seedings for each matrix type. (C) Representative widefield images of cropped channel ROIs showing GFP^+^-KP4 cells in Matrigel, Rat Tail Collagen Type I, and Bovine Collagen Type I in thermoformed 96- GLAnCE immediately after seeding (Day 0) and on Days 2, 4, and 6. Scale bars are 500 µm.

### 3.5 Application of the thermoformed 96-GLAnCE for a longitudinal image-based cell proliferation assay

Confident that the manufacturing and seeding of thermoformed chips were sufficiently robust, we next sought to use our thermoformed 96-GLAnCE platform for a longitudinal image-based cell proliferation assay using the pancreatic cancer cell line KP4, transduced with a green fluorescent protein (GFP^+^ KP4). Previously, we observed that the growth of this cell line could not be easily tracked over time in the original 96-GLAnCE due to its highly proliferative and contractile behavior, leading to microgel detachment. Since the non-vertical channel walls slightly increased the channel wall area in thermoformed chips, we hypothesized that the increased surface area could improve gel binding to the polystyrene surface and resist cell-induced gel contraction over longer experimental timelines. When we cultured GFP^+^ KP4 cells in the standard Rat Tail Collagen matrix in the thermoformed chips, we were able to monitor cell proliferation over 6 days while observing minimal gel contraction (after which experiments were stopped due to cells proliferating outside microgels) (Figure 6A, B).

**Figure 6.**
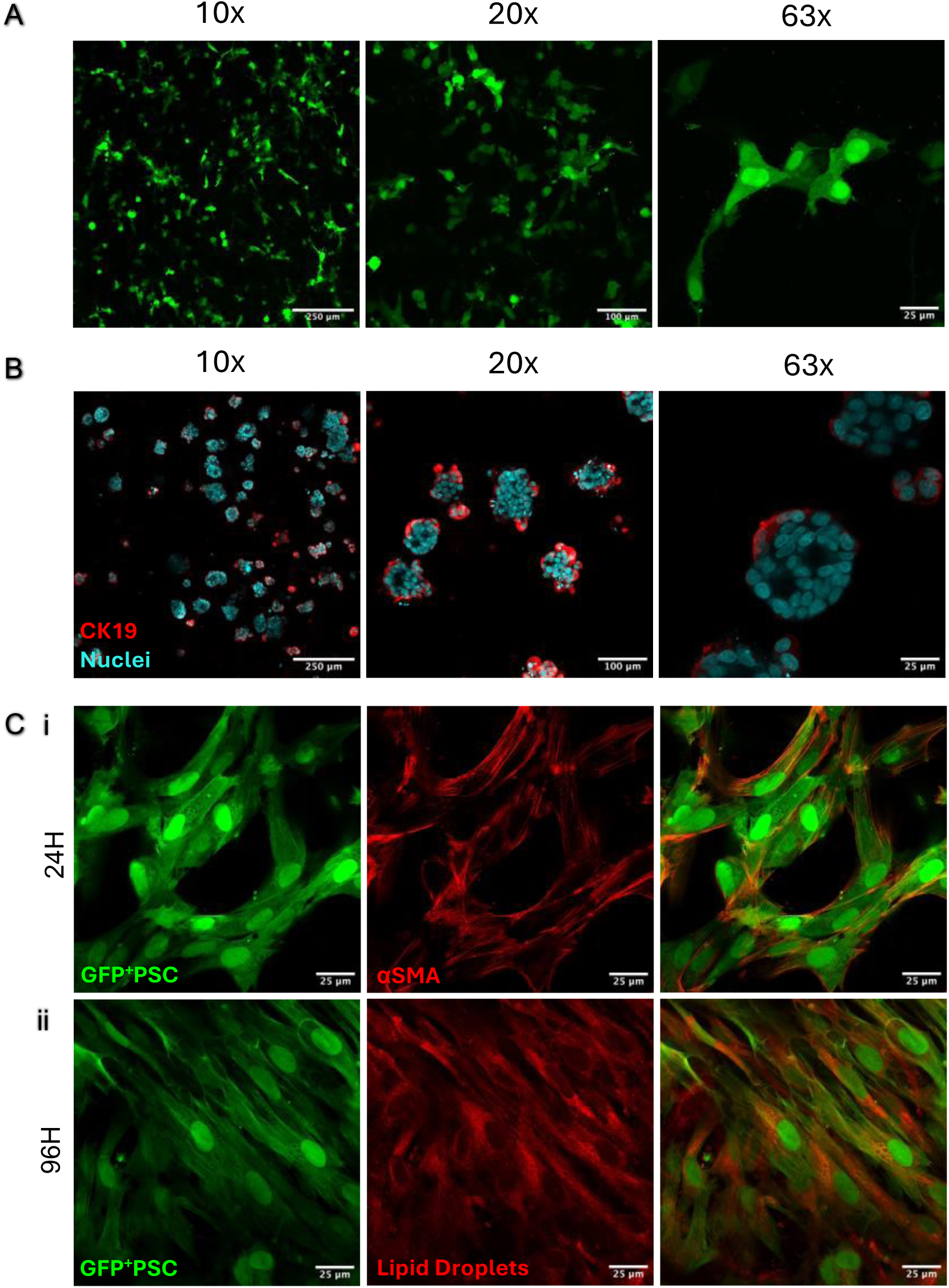
Assessing compatibility of thermoformed 96-GLAnCE with confocal microscopy and imaging at high magnifications. (A) Representative confocal images of GFP^+^ KP4 cells (green) in thermoformed 96-GLAnCE after 3 days of culture. Images were acquired at 10x, 20x, and 63x on Leica SP8. Scale bars are 250, 100, and 25 µm, respectively. (B) Representative confocal images of pancreatic cancer patient-derived organoids (PPTO.46) that were immuno-fluorescently stained for Cytokeratin 19 (CK19, red) and Nuclei (DRAQ5, cyan) after 5 days of culture. Images were acquired at 10x, 20x, and 63x on Leica SP8. Scale bars are 250, 100, and 25 µm, respectively. (C) Representative confocal images of GFP^+^ pancreatic stellate cells (GFP^+^ PSCs, green) that were immuno-fluorescently stained for (i) alpha-smooth muscle actin (αSMA, red) and (ii) lipid droplets (BODIPY Red, red) after 1 and 4 days of culture, respectively. Images were acquired at 63x on Leica SP8. Scale bars are 25 µm.

Given the feasibility of culturing the GFP^+^ KP4 cells in the thermoformed chip over time, we also tested other commonly used natural protein matrices, Matrigel and Bovine Collagen Type I. While we previously reported that the original chips were compatible with Matrigel (25), we did observe higher rates of microgel damage during the removal of the PDMS slab after gel seeding. However, we observed only minor damage to Matrigel-based microgels in thermoformed chips, potentially due to the sloping channel wall profiles. This showcases that our thermoformed device offers improved capabilities over the original chips beyond just logistics.

We could also track cell proliferation in Matrigel and Bovine collagen for up to 1 week (Figure 6B); however, we did notice more extensive gel shrinkage at channel ends (Supplementary Figure 6Ai-iii). The shrinkage could be due to the physical properties of the matrix, as the Rat Tail Collagen forms shorter fibrils and smaller pores when used at the same concentration, thus yielding much denser fiber networks that take more time to remodel (46). In turn, the intrinsic capacity of GFP^+^ KP4 cancer cells to remodel the matrix may also differ depending on their environment. Interestingly, we also observed different GFP^+^ KP4 morphologies and proliferation patterns for all three matrices by Day 4 (Figure 5C). Cells acquired a spread-out morphology in Matrigel and Rat Tail Collagen but remained circular in Bovine Collagen. Cells appeared to grow in clusters unevenly throughout the gel in Matrigel, whereas more homogeneous cell distribution was observed for Rat Tail and Bovine collagens. Although these observations for GFP^+^-KP4 cells require further investigation, this preliminary work highlights the significance of imaging readouts in ECM-based models for providing more granular insights about cell phenotypic changes and behavior in different 3D environments compared to standard soluble bulk assays.

In this study, as a proof-of-concept, we performed a longitudinal proliferation assay with the cancer cell line cultured alone. In the future, however, tumor stromal cells such as fibroblasts and immune cells could also be incorporated into thermoformed 96-GLAnCE cultures, given the robustness of ECM gels to remain intact and immobilized over extended culture periods. Given many of these stromal cells perform localized behaviors in response to contact with cancer cells, we envision the thermoformed 96-GLAnCE will enable the development of stromal cell-centered imaging-based assays such as migration/ invasion, phagocytosis, and ECM deposition and alignment (10,47–49); the platform could additionally implement more sophisticated fluorescent reporter systems (50,51) for mechanistic studies of these cellular processes.

### 3.6 Application of the thermoformed 96-GLAnCE for imaging using confocal microscopy

Confident that the thermoformed chips enabled longitudinal imaging using widefield fluorescence microscopy, we next wanted to apply the thermoformed 96-GLAnCE in imaging-based readouts using confocal microscopy. The vacuum thermoforming fabrication method allowed for a dramatic reduction in the thickness of the 96-GLAnCE bottom component (fourteen-fold), thus rendering the platform compatible with high-magnification objectives. This significant improvement allowed us to image GFP^+^ KP4 cells in the thermoformed devices using 10x (NA 0.4), 20x (NA 0.75), and 63x (NA 1.40) objectives (Figure 6A). At 63x magnification, we could distinguish intracellular features and observed a cell undergoing a nuclear division. The detail resolution and image quality were comparable to the standard 2D glass coverslips (Supplementary Figure 7).

Building on our work using cell lines, we also incorporated primary pancreatic ductal adenocarcinoma patient-derived organoids (PDAC PDOs) into the thermoformed 96-GLAnCE and performed *in situ* immunofluorescence staining for an epithelial marker Cytokeratin 19 (CK19) and nuclei. We could robustly visualize cells at 10x, 20x, and 63x magnifications. PDAC PDOs preserved their epithelial-like phenotype and polarized morphology in the thermoformed 96-GLAnCE, as evident from CK19 expression and lumens in 63x images. To confirm that other intracellular features of primary cells could be visualized at 63x magnification, we also cultured primary pancreatic stellate cells, expressing GFP (GFP^+^ PSCs) in the thermoformed 96-GLAnCE for 24 and 96 hours and stained for the cytoskeletal protein alpha-smooth muscle actin (αSMA) (Figure 6Ci) and lipid droplets (Figure 6Cii), respectively. We selected these markers because they allow monitoring of PSC phenotypic plasticity in response to the substrate stiffness. Specifically, when PSCs transition into a quiescent-like phenotype when cultured in 3D gel environments as opposed to stiff tissue culture plastic, cells show a spread-out morphology and accumulate lipid droplets over time (52,53). In the 96- GLAnCE chips, we observed that GFP^+^ PSCs had a spindle-like morphology and prominently expressed the activation marker aSMA at 24 hours. In contrast, by 96 hours, GFP^+^ PSCs had acquired a spread-out morphology and clusters of lipid droplets in their cytoplasm.

These qualitative observations demonstrate that thermoformed 96-GLAnCE is a versatile platform capable of supporting cell lines, primary cells, and organoids. It is also compatible with confocal microscopy and high-magnification objectives, eliminating the need for sample pre-processing or optical clearing steps. By combining this platform with high-throughput confocal microscopy, researchers can study cell behavior at single-cell and subcellular resolution, enabling the detection of regulatory proteins with low expression levels that would otherwise be undetectable at lower magnifications. Currently, single-cell analysis of organoids remains challenging due to the compact density of cells within their structure. However, with the growing use of Artificial Intelligence and deep learning models, image analysis pipelines for effective organoid single-cell segmentation are emerging (54,55). In this context, the high- quality and high-resolution confocal images obtained from the thermoformed 96-GLAnCE will provide valuable datasets as more advanced segmentation and analysis workflows continue to develop.

## 4. Conclusions

In this report, we developed an accessible approach to fabricate 96-GLAnCE chips from thin polystyrene films using vacuum thermoforming to replace previous manufacturing from thick polystyrene sheets by hot-embossing. We demonstrated we could thermoform an array of mm- scale channels using a commercially available desktop thermoformer, a shop vacuum (for pressure differential), and a custom positive aluminum mold. Our final operating conditions allowed reductions in both cost (50% reduction) and fabrication time (40% reduction) compared to the hot-embossing method used for the original 96-GLAnCE manufacturing. We showed that thermoformed chips were compatible with existing seeding and device assembly workflows and could be used for a longitudinal imaging-based assay in different seeding hydrogel matrices with minimal gel contraction after 1 week of cell culture. These experiments highlighted the flexibility of our device for asking different biological questions requiring analysis over various time scales. Notably, the vacuum thermoforming fabrication method allowed for a dramatic reduction in the thickness of the 96-GLAnCE chips (fourteen-fold), thus rendering the platform compatible with the working distance of high magnification objectives and allowing for image acquisition at single-cell and subcellular resolution using confocal microscopy. Combining our device with high-throughput confocal microscopy could provide a useful platform for studying important regulatory proteins with low expression levels that are otherwise not detectable at low magnifications. We envision that the ongoing improvements in effective image segmentation and analysis pipelines will enable the analysis of individual cells within the compact density of organoid structures using images like those obtained with our thermoformed chips.

## 5. Declarations

### 5.1 Abbreviations

αSMA: Alpha Smooth Muscle Actin
BF: Brightfield
CK19: Cytokeratin 19
CV: Coefficient of Variation
DAPI: 4’,6-Diamidino-2-Phenylindole
DMEM: Dulbecco’s Modified Eagle Medium
ECM: Extracellular Matrix
FBS: Fetal Bovine Serum
FITC: Fluorescein Isothiocyanate
GLAnCE: Gels for Live Analysis of Compartmentalized Environments
GFP: Green Fluorescent Protein
IMDM: Iscove’s Modified Dulbecco’s Medium
MGV: Mean Gray Value
PBS: Phosphate-Buffered Saline
PDMS: Polydimethylsiloxane
PDO: Patient-Derived Organoid
PDAC: Pancreatic Ductal Adenocarcinoma
PSC: Pancreatic Stellate Cell
PS: Polystyrene
ROI: Region of Interest

### 5.2 Ethics approval and consent to participants

Pancreatic ductal adenocarcinoma organoids (PMLB identifier PPTO.46 from the University Health Network, Ontario, Canada) were obtained from the UHN Biobank at Princess Margaret Cancer Center in compliance with the University of Toronto Research Ethics Board guidelines (protocol #36107).

### 5.3 Consent for publication

Not applicable.

### 5.4 Availability of data and materials

The data supporting this article have been deposited and are available at: McGuigan, Alison (2024), “Fomina et al 2024”, Mendeley Data, V1, doi: 10.17632/m4mhd788hp.1

### 5.7 Authors’contributions

AF, NCW, and NTL conceived of the project. AF designed and performed experiments, analyzed data, and prepared figures with assistance from AT and JL. APM supervised and funded the research and designed experiments. AF and APM wrote the manuscript. All authors edited and approved the manuscript.

## Acknowledgements

Schematics were prepared using BioRender.com and Vectary.com.

## References

1. Sung H, Ferlay J, Siegel RL, Laversanne M, Soerjomataram I, Jemal A, et al. Global Cancer Statistics 2020: GLOBOCAN Estimates of Incidence and Mortality Worldwide for 36 Cancers in 185 Countries. CA Cancer J Clin. 2021 May;71(3):209–49.

2. Hanahan D. Hallmarks of Cancer: New Dimensions. Cancer Discov. 2022;12(1):31–46.

3. Horvath P, Aulner N, Bickle M, Davies AM, Nery ED, Ebner D, et al. Screening out irrelevant cell-based models of disease. Nat Rev Drug Discov. 2016 Nov;15(11):751–69.

4. Rodenhizer D, Dean T, D’Arcangelo E, McGuigan AP. The Current Landscape of 3D In Vitro Tumor Models: What Cancer Hallmarks Are Accessible for Drug Discovery? Adv Healthc Mater. 2018 Apr;7(8):1701174.

5. Kessel S, Cribbes S, Déry O, Kuksin D, Sincoff E, Qiu J, et al. High-Throughput 3D Tumor Spheroid Screening Method for Cancer Drug Discovery Using Celigo Image Cytometry. SLAS Technol. 2017 Aug;22(4):454–65.

6. Jenkins RW, Aref AR, Lizotte PH, Ivanova E, Stinson S, Zhou CW, et al. Ex Vivo Profiling of PD-1 Blockade Using Organotypic Tumor Spheroids. Cancer Discov. 2018 Feb;8(2):196–215.

7. Bauleth-Ramos T, Feijão T, Gonçalves A, Shahbazi MA, Liu Z, Barrias C, et al. Colorectal cancer triple co-culture spheroid model to assess the biocompatibility and anticancer properties of polymeric nanoparticles. J Controlled Release. 2020 Jul;323:398–411.

8. Schuster B, Junkin M, Kashaf SS, Romero-Calvo I, Kirby K, Matthews J, et al. Automated microfluidic platform for dynamic and combinatorial drug screening of tumor organoids. Nat Commun. 2020 Oct 19;11(1):5271.

9. Geyer M, Schreyer D, Gaul LM, Pfeffer S, Pilarsky C, Queiroz K. A microfluidic-based PDAC organoid system reveals the impact of hypoxia in response to treatment. Cell Death Discov. 2023 Jan 21;9(1):20.

10. Heaster TM, Humayun M, Yu J, Beebe DJ, Skala MC. Autofluorescence Imaging of 3D Tumor–Macrophage Microscale Cultures Resolves Spatial and Temporal Dynamics of Macrophage Metabolism. Cancer Res. 2020;80(23):5408–23.

11. Landon-Brace N, Cadavid JL, Latour S, Co IL, Rodenhizer D, Li NT, et al. An Engineered Patient-Derived Tumor Organoid Model That Can Be Disassembled to Study Cellular Responses in a Graded 3D Microenvironment. Adv Funct Mater. 2021;2105349.

12. Li NT, Wu NC, Cao R, Cadavid JL, Latour S, Lu X, et al. An off-the-shelf multi-well scaffold-supported platform for tumour organoid-based tissues. Biomaterials. 2022 Dec;291:121883.

13. Tang M, Xie Q, Gimple RC, Zhong Z, Tam T, Tian J, et al. Three-dimensional bioprinted glioblastoma microenvironments model cellular dependencies and immune interactions. Cell Res. 2020 Oct;30(10):833–53.

14. Pierrevelcin M, Flacher V, Mueller CG, Vauchelles R, Guerin E, Lhermitte B, et al. Engineering Novel 3D Models to Recreate High-Grade Osteosarcoma and its Immune and Extracellular Matrix Microenvironment. Adv Healthc Mater. 2022 Oct;11(19):2200195.

15. Flores-Torres S, Dimitriou NM, Pardo LA, Kort-Mascort J, Pal S, Peza-Chavez O, et al. Bioprinted Multicomponent Hydrogel Co-culture Tumor-Immune Model for Assessing and Simulating Tumor-Infiltrated Lymphocyte Migration and Functional Activation. ACS Appl Mater Interfaces. 2023 Jul 19;15(28):33250–62.

16. Drost J, Clevers H. Organoids in cancer research. Nat Rev Cancer. 2018 Jul;18(7):407– 18.

17. Tuveson D, Clevers H. Cancer modeling meets human organoid technology. Science. 2019 Jun 7;364(6444):952–5.

18. Phan N, Hong JJ, Tofig B, Mapua M, Elashoff D, Moatamed NA, et al. A simple high- throughput approach identifies actionable drug sensitivities in patient-derived tumor organoids. Commun Biol. 2019 Dec;2(1):78.

19. Hou S, Tiriac H, Sridharan BP, Scampavia L, Madoux F, Seldin J, et al. Advanced Development of Primary Pancreatic Organoid Tumor Models for High-Throughput Phenotypic Drug Screening. SLAS Discov Adv Sci Drug Discov. 2018 Jul;23(6):574–84.

20. Dijkstra KK, Cattaneo CM, Weeber F, Chalabi M, Van De Haar J, Fanchi LF, et al. Generation of Tumor-Reactive T Cells by Co-culture of Peripheral Blood Lymphocytes and Tumor Organoids. Cell. 2018;174(6):1586–1598.e12.

21. Dekkers JF, Alieva M, Cleven A, Keramati F, Wezenaar AKL, van Vliet EJ, et al. Uncovering the mode of action of engineered T cells in patient cancer organoids. Nat Biotechnol. 2023 Jan;41(1):60–9.

22. Neal JT, Li X, Zhu J, Giangarra V, Grzeskowiak CL, Ju J, et al. Organoid Modeling of the Tumor Immune Microenvironment. Cell. 2018 Dec;175(7):1972–1988.e16.

23. Jacob F, Salinas RD, Zhang DY, Nguyen PTT, Schnoll JG, Wong SZH, et al. A Patient- Derived Glioblastoma Organoid Model and Biobank Recapitulates Inter- and Intra- tumoral Heterogeneity. Cell. 2020 Jan 9;180(1):188–204.e22.

24. Lin S, Schorpp K, Rothenaigner I, Hadian K. Image-based high-content screening in drug discovery. Drug Discov Today. 2020 Aug;25(8):1348–61.

25. D’Arcangelo E, Wu NC, Chen T, Shahaj A, Cadavid JL, Huang L, et al. Gels for Live Analysis of Compartmentalized Environments (GLAnCE): A tissue model to probe tumour phenotypes at tumour-stroma interfaces. Biomaterials. 2020 Jan;228:119572.

26. Wu NC, Cadavid JL, Tan X, Latour S, Scaini S, Makhijani P, et al. 3D microgels to quantify tumor cell properties and therapy response dynamics. Biomaterials. 2022 Apr;283:121417.

27. Zhang Y, Gross H. Systematic design of microscope objectives. Part I: System review and analysis. Adv Opt Technol. 2019 Aug 27;8(5):313–47.

28. Throne J. Thermoforming. In: Applied Plastics Engineering Handbook [Internet]. Elsevier; 2017 [cited 2024 Sep 12]. p. 345–75. Available from: https://linkinghub.elsevier.com/retrieve/pii/B978032339040800016X

29. Vrij EJ, Espinoza S, Heilig M, Kolew A, Schneider M, Van Blitterswijk CA, et al. 3D high throughput screening and profiling of embryoid bodies in thermoformed microwell plates. Lab Chip. 2016;16(4):734–42.

30. Giselbrecht S, Gietzelt T, Gottwald E, Trautmann C, Truckenmüller R, Weibezahn KF, et al. 3D tissue culture substrates produced by microthermoforming of pre-processed polymer films. Biomed Microdevices. 2006;8(3):191–9.

31. Fois MG, Tahmasebi Birgani ZN, López-Iglesias C, Knoops K, van Blitterswijk C, Giselbrecht S, et al. In vitro vascularization of 3D cell aggregates in microwells with integrated vascular beds. Mater Today Bio. 2024 Dec;29:101260.

32. Iggo R, Richard E. Lentiviral Transduction of Mammary Epithelial Cells. In: Vivanco MDM, editor. Mammary Stem Cells [Internet]. New York, NY: Springer New York; 2015 [cited 2024 Sep 12]. p. 137–60. (Methods in Molecular Biology; vol. 1293). Available from: https://link.springer.com/10.1007/978-1-4939-2519-3_8

33. Preibisch S, Saalfeld S, Tomancak P. Globally optimal stitching of tiled 3D microscopic image acquisitions. Bioinformatics. 2009 Jun 1;25(11):1463–5.

34. Riley A. Plastics manufacturing processes for packaging materials. In: Packaging Technology [Internet]. Elsevier; 2012 [cited 2024 Apr 2]. p. 310–60. Available from: https://linkinghub.elsevier.com/retrieve/pii/B9781845696658500147

35. Nguyen TK, Lee BK. Investigation of processing parameters in micro-thermoforming of micro-structured polystyrene film. J Mech Sci Technol. 2019 Dec;33(12):5669–75.

36. Wypych G. PS polystyrene. In: Handbook of Polymers [Internet]. Elsevier; 2012 [cited 2024 Sep 12]. p. 541–7. Available from: https://linkinghub.elsevier.com/retrieve/pii/B9781895198478501624

37. Sehrawat M, Rani M, Bharadwaj S, Sharma S, Chauhan GS, Dhakate SR, et al. Glass Transition Temperature Measurement of Polycarbonate Specimen by Dynamic Mechanical Analyser Towards the Development of Reference Material. MAPAN. 2022 Sep;37(3):517–27.

38. Hajduk B, Bednarski H, Jarka P, Janeczek H, Godzierz M, Tański T. Thermal and optical properties of PMMA films reinforced with Nb2O5 nanoparticles. Sci Rep. 2021 Nov 18;11(1):22531.

39. Agha A, Waheed W, Alamoodi N, Mathew B, Alnaimat F, Abu-Nada E, et al. A Review of Cyclic Olefin Copolymer Applications in Microfluidics and Microdevices. Macromol Mater Eng. 2022 Aug;307(8):2200053.

40. Truckenmüller R, Giselbrecht S, Van Blitterswijk C, Dambrowsky N, Gottwald E, Mappes T, et al. Flexible fluidic microchips based on thermoformed and locally modified thin polymer films. Lab Chip. 2008;8(9):1570.

41. Truckenmüller R, Giselbrecht S, Rivron N, Gottwald E, Saile V, van den Berg A, et al. Thermoforming of film-based biomedical microdevices. Adv Mater Deerfield Beach Fla. 2011 Mar 18;23(11):1311–29.

42. Jain N, Vogel V. Spatial confinement downsizes the inflammatory response of macrophages. Nat Mater. 2018 Dec;17(12):1134–44.

43. Nikolaev M, Mitrofanova O, Broguiere N, Geraldo S, Dutta D, Tabata Y, et al. Homeostatic mini-intestines through scaffold-guided organoid morphogenesis. Nature. 2020 Sep 24;585(7826):574–8.

44. Korolj A, Laschinger C, James C, Hu E, Velikonja C, Smith N, et al. Curvature facilitates podocyte culture in a biomimetic platform. Lab Chip. 2018;18(20):3112–28.

45. Samal P, Gubbins E, Van Blitterswijk C, Truckenmüller R, Giselbrecht S. Thin fluorinated polymer film microcavity arrays for 3D cell culture and label-free automated feature extraction. Biomater Sci. 2021;9(23):7838–50.

46. Wolf K, Alexander S, Schacht V, Coussens LM, Von Andrian UH, Van Rheenen J, et al. Collagen-based cell migration models in vitro and in vivo. Semin Cell Dev Biol. 2009;20(8):931–41.

47. Parlato S, De Ninno A, Molfetta R, Toschi E, Salerno D, Mencattini A, et al. 3D Microfluidic model for evaluating immunotherapy efficacy by tracking dendritic cell behaviour toward tumor cells. Sci Rep. 2017 Dec;7(1):1093.

48. Martinez-Marin D, Jarvis C, Nelius T, De Riese W, Volpert OV, Filleur S. PEDF increases the tumoricidal activity of macrophages towards prostate cancer cells in vitro. Tang CH, editor. PLOS ONE. 2017 Apr 12;12(4):e0174968.

49. Jones CE, Sharick JT, Sizemore ST, Cukierman E, Strohecker AM, Leight JL. A Miniaturized Screening Platform to Identify Novel Regulators of Extracellular Matrix Alignment. Cancer Res Commun. 2022 Nov 22;2(11):1471–86.

50. Burgstaller S, Bischof H, Gensch T, Stryeck S, Gottschalk B, Ramadani-Muja J, et al. pH-Lemon, a Fluorescent Protein-Based pH Reporter for Acidic Compartments. ACS Sens. 2019;4(4):883–91.

51. Sung BH, Von Lersner A, Guerrero J, Krystofiak ES, Inman D, Pelletier R, et al. A live cell reporter of exosome secretion and uptake reveals pathfinding behavior of migrating cells. Nat Commun. 2020 Apr 29;11(1):2092.

52. Jesnowski R, Fürst D, Ringel J, Chen Y, Schrödel A, Kleeff J, et al. Immortalization of pancreatic stellate cells as an in vitro model of pancreatic fibrosis: deactivation is induced by matrigel and N-acetylcysteine. Lab Invest. 2005 Oct;85(10):1276–91.

53. Lachowski D, Cortes E, Pink D, Chronopoulos A, Karim SA, P. Morton J, et al. Substrate Rigidity Controls Activation and Durotaxis in Pancreatic Stellate Cells. Sci Rep. 2017 May 31;7(1):2506.

54. Mukashyaka P, Kumar P, Mellert DJ, Nicholas S, Noorbakhsh J, Brugiolo M, et al. High-throughput deconvolution of 3D organoid dynamics at cellular resolution for cancer pharmacology with Cellos. Nat Commun. 2023 Dec 18;14(1):8406.

55. Liu B, Zhu Y, Yang Z, Yan HHN, Leung SY, Shi J. Deep Learning–Based 3D Single-Cell Imaging Analysis Pipeline Enables Quantification of Cell–Cell Interaction Dynamics in the Tumor Microenvironment. Cancer Res. 2024 Feb 15;84(4):517–26.

