## Supplementary information for "In-House Manufacturing of 3D Culture Chips via Vacuum Thermoforming for Enhanced Imaging Applications"

**Vacuum Thermoforming for the Fabrication of *In Vitro* Culture Chips: Enabling High-Resolution Imaging in 3D**

**Supplemental figure captions.**

**Supplementary Figure 1. Desktop thermoformer properties and PDMS casting workflow.** (A) Overview of Mayuku Formbox desktop thermoformer. The vacuum bed holds a custom mold, allowing vacuum suction across the mold surface. Trays allow for securing a sheet of plastic film and bringing it up close to the heating element, which heats the plastic material to a malleable state. The thermoformer has 6 temperature settings. Diagrams created with Vectary.com. (B) Thermoformer temperature settings and their corresponding absolute temperatures as manually measured using a thermometer after running each setting for 10 minutes. The measurements were repeated 3 times for each temperature setting. Thermoformer temperature settings and absolute temperatures show linear relationship ($R^2 = 0.9916$) (C) Overview of the PDMS casting workflow used to characterize vacuum thermoformed samples. PDMS solution was poured into vacuum thermoformed samples and cured overnight. Casted samples were cut and bisected to obtain cross-sections. Sample cross-sections were imaged using a brightfield microscope; sample features were quantified from brightfield images in FIJI. Diagrams created with Biorender.com.

**Supplementary Figure 2. Selecting mold vacuum hole configuration and preheating temperature using a 3 × 3 prototype for thermoforming 96-GLAnCE chips.** (A) Design of a custom positive aluminum mold with a 3 × 3 array of microchannels and nine vacuum hole configurations (labeled from 1 to 9). For specific dimensions of each vacuum hole configuration, please refer to Table 1. Diagrams created with Vectary.com. (B) (i) A representative 4X brightfield image of a microchannel cross-section used to quantify channel angle after thermoforming. Scale bar is 500 μm. (ii) Quantification of channel angle of thermoformed channels after the 3 × 3 mold preheating at 120 and 150 °C in the oven. Each plotted point represents an independent sample. Statistics performed using unpaired t-test using

Welch's correction, ***p< 0.001. N = 3-4 independent samples. (iii) Quantification of channel angle for thermoformed channels 7, 8, and 9 when mold was preheated at 150 °C in the oven. Each plotted point represents an independent sample. Statistics performed using ordinary one-way ANOVA, ns – no significance. N = 3 independent film samples. (iv) Summary of mean angle values ± SD for channels 7, 8, and 9 when mold was preheated at 150 °C in the oven.

**Supplementary Figure 3. Channel depth comparison between thermoformed and original 96-GLAnCE chips.** Quantification of channel depth of thermoformed and original hot-embossed 96-GLAnCE chips. Each plotted point represents an independent chip. Statistics performed using unpaired t-test using Welch's correction, ns – no significance. N = 3-4 independent chips.

**Supplementary Figure 4. Characterization of autofluorescence of original and thermoformed 96-GLAnCE chips.** (A) Representative 4x fluorescence images showing autofluorescence of original (top) and thermoformed 96-GLAnCE (bottom) in the DAPI (blue), FITC (green), Cy3 (yellow), Texas Red (red), and Cy5 (magenta) channels. Each fluorescent image was taken with an exposure time of 300 ms. Scale bar is 500 μm. (B) Quantification of autofluorescence based on mean gray value (MGV) for each fluorescence channel and type of chip. Each plotted point represents an independent chip. Statistics performed using unpaired t-test with Welch's correction; **, p< 0.01; ****, p< 0.0001. N = 4 independent original and thermoformed 96-GLAnCE chips.

**Supplementary Figure 5. Selecting gel volume for thermoformed 96-GLAnCE seeding.** (A) (i) Representative widefield images of thermoformed 96-GLAnCE microchannels (2 sites) seeded with 3.0, 3.3, and 3.6 μL of Rat Tail collagen Type I solution (3mg/mL) containing fluorescent FITC beads. Scale bar is 1000 μm. (ii) Quantification of % channel area occupied by fluorescent FITC beads for each gel volume tested. Each plotted point represents a seeded channel. Statistics performed using ordinary one-way ANOVA; **, p< 0.01 ****, p< 0.0001. N = 3 independent seedings. (B) (i) Representative widefield image of a thermoformed 96-GLAnCE microchannel (1 site) seeded with 3.6 μl of Rat Tail collagen Type I solution (3mg/mL) containing fluorescent FITC beads and (ii) its cropped region of interest (ROI, 1100x700 pixels). Scale bar is 1000 μm. (B) (i) Representative widefield image of an original 96-GLAnCE microchannel (1 site) seeded with 3.0 μl of Rat Tail collagen Type I solution

(3mg/mL) containing fluorescent FITC beads and (ii) its cropped region of interest (ROI, 1100x700 pixels). Scale bar is 1000 μm.

**Supplementary Figure 6. Gel contraction in different matrices in thermoformed 96-GLAnCE.** Representative widefield images (2 sites) of GFP-KP4 cells in Rat Tail Collagen Type I, Matrigel, and Bovine Collagen Type I on Day 6 after seeding in a thermoformed 96-GLAnCE chip. Scale bar is 1000 μm.

**Supplementary Figure 7. Assessing compatibility of thermoformed 96-GLAnCE with confocal microscopy and imaging at high magnifications.** Representative confocal images of $GFP^+$ KP4 cells (green) on glass coverslips after 3 days of culture. Images were acquired at 10X, 20X, and 63X on Leica SP8. Scale bars are 250, 100, and 25 μm, respectively.

**Supplementary Figure 1.**

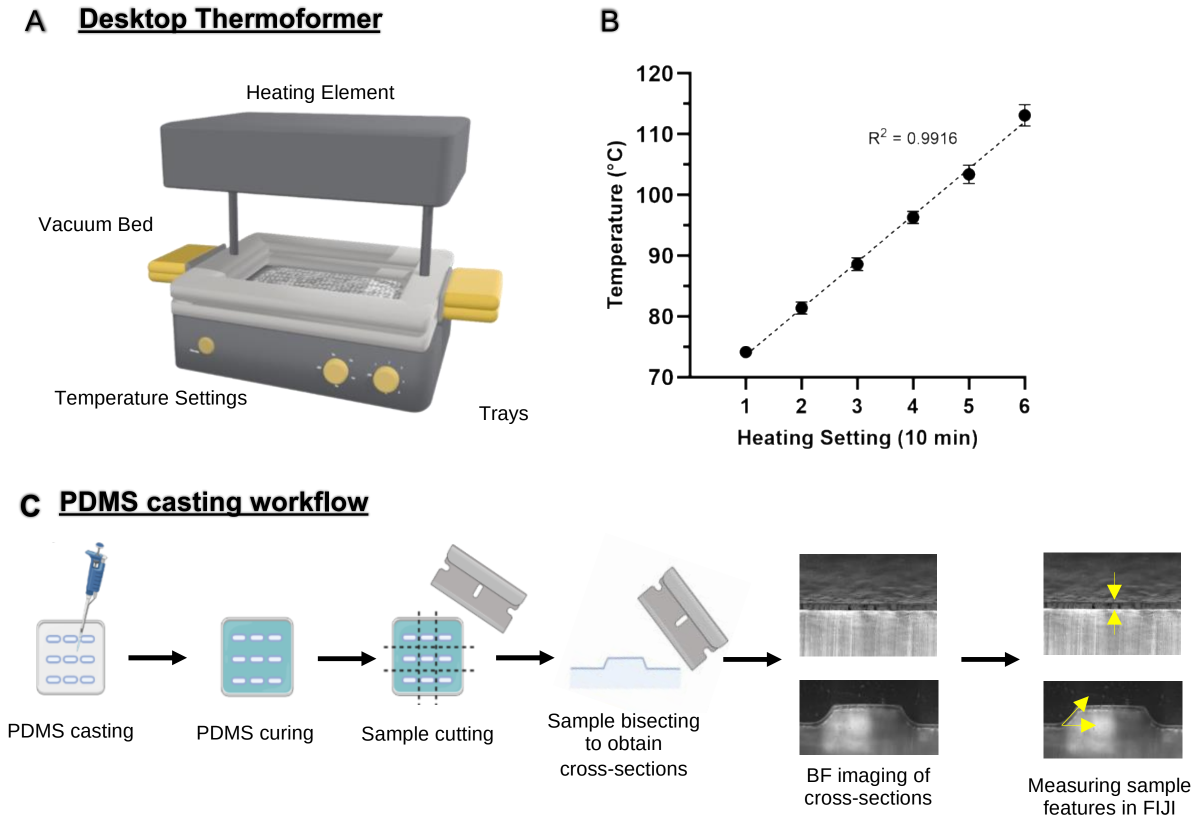

**Supplementary Figure 2.**

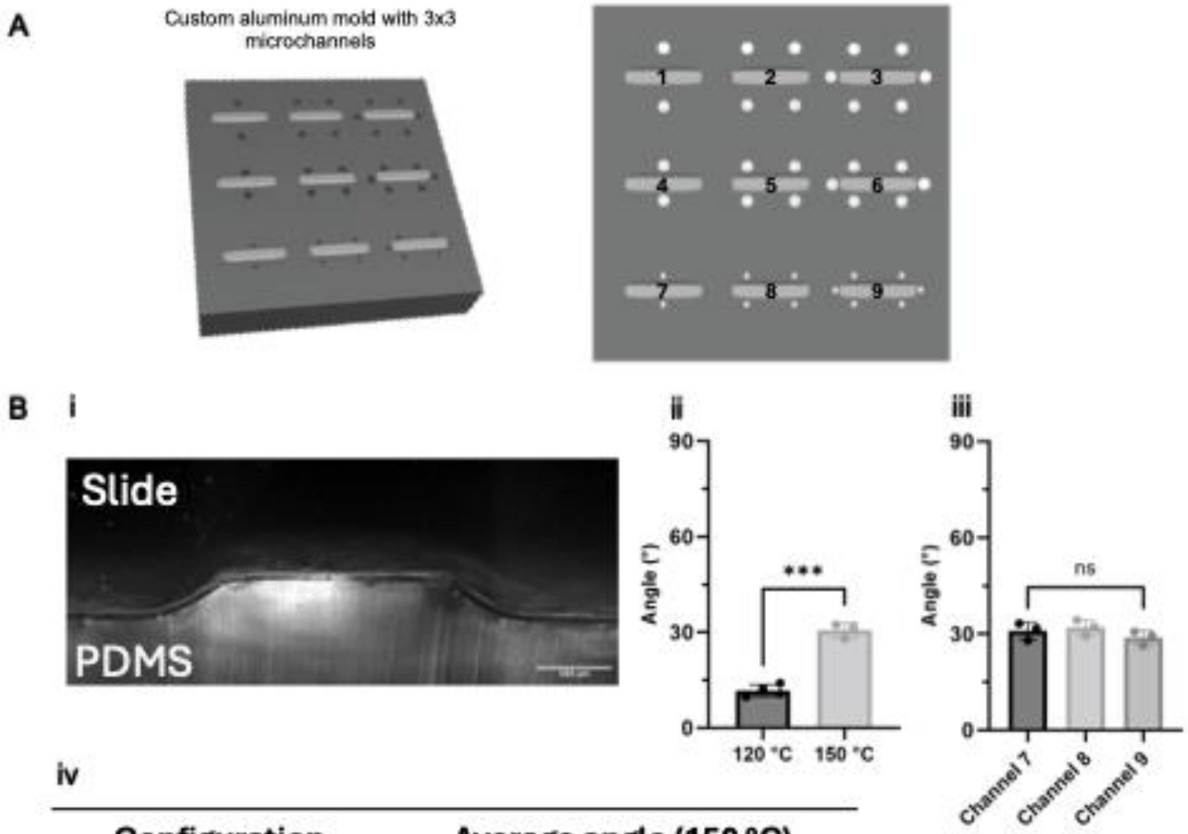


iv

| Configuration | Average angle (150 °C) |
|---|---|
| Channel 7 | 31.0 ± 2.7 |
| Channel 8 | 32.0 ± 2.5 |
| Channel 9 | 28.9 ± 2.3 |

**Supplementary Figure 3.**

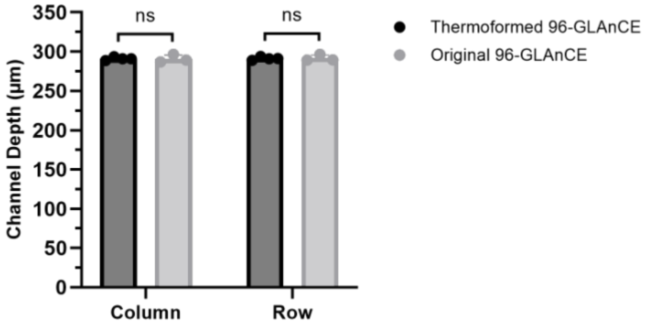

### Supplementary Figure 4.

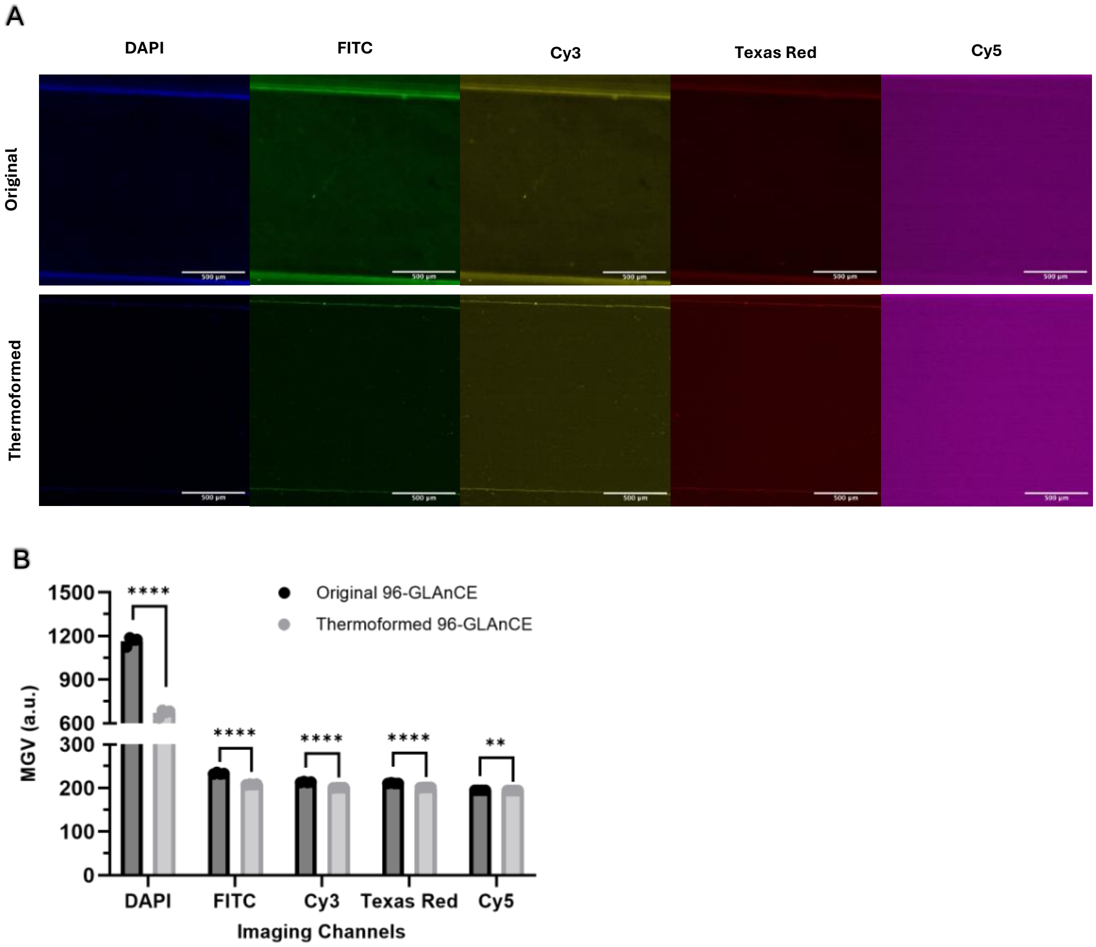

### Supplementary Figure 5.

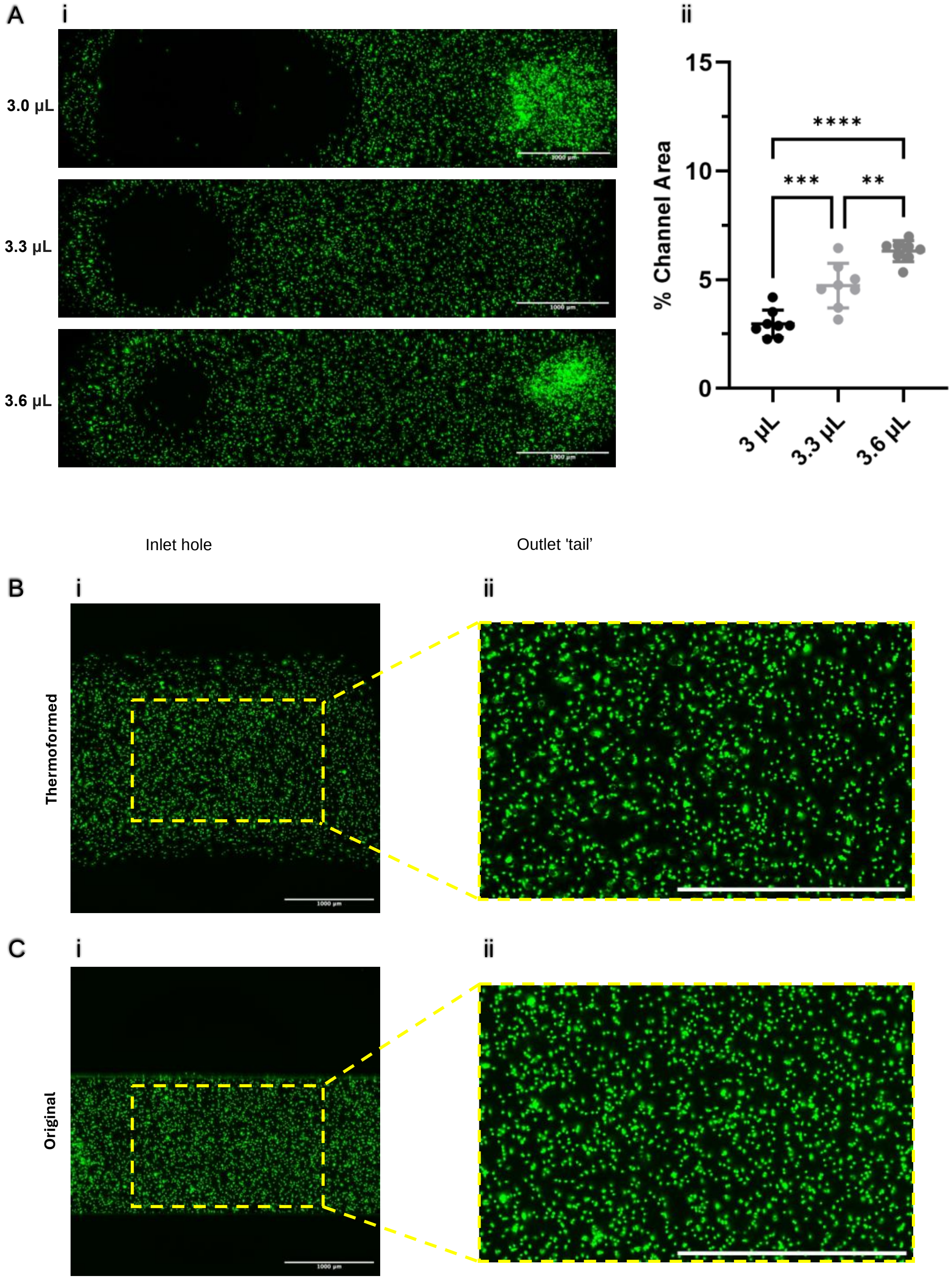

**Supplementary Figure 6.**

Day 6

Rat Tail Type I Collagen

Matrigel

Bovine Type I Collagen

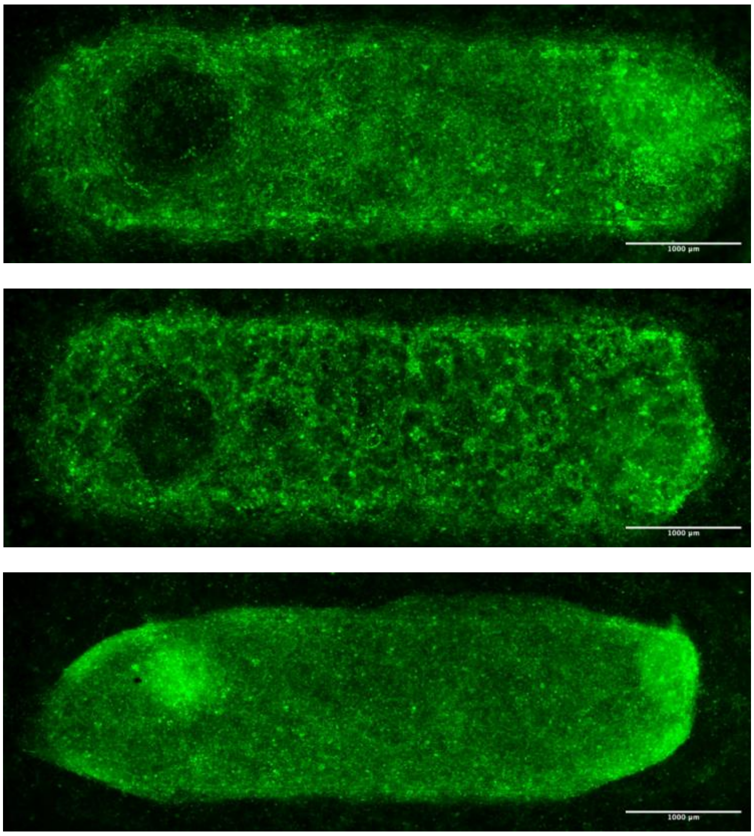

**Supplementary Figure 7.**

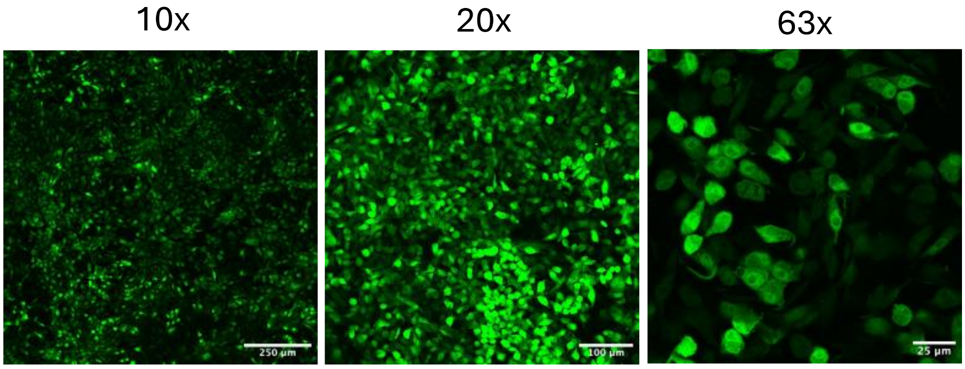
